# An auxiliary protein tunes reductase activity in alkaloid biosynthesis

**DOI:** 10.64898/2026.04.21.719811

**Authors:** Samuel C. Carr, Song Wu, Klaus Gase, Yoko Nakamura, Mohamed Omar Kamileen, Maritta Kunert, Sarah Heinicke, Moonyoung Kang, Veit Grabe, Lorenzo Caputi, Sarah E. O’Connor

**Affiliations:** Department of Natural Product Biosynthesis, Max Planck Institute for Chemical Ecology, 07745 Jena, Germany; Microscopic Imaging Service Group, Max Planck Institute for Chemical Ecology, 07745 Jena, Germany

**Keywords:** Natural Products, Biosynthesis, Alkaloids, Auxiliary Proteins

## Abstract

Monoterpene indole alkaloids encompass a diverse class of plant natural products. Many of these valuable compounds, including the anticancer medicines vinblastine and vincristine (*Catharanthus roseus*) and the antiaddiction treatment ibogaine (*Tabernanthe iboga)* utilize a common biosynthetic intermediate, 19*E*-geissoschizine, which is produced by the medium-chain dehydrogenase/reductase geissoschizine synthase (GS). Herein we report the discovery of a non-catalytic auxiliary protein (*facilitator of geissoschizine synthase*, FoGS) that improves formation of 19*E*-geissoschizine by acting in combination with GS. The discovery of a FoGS orthologue that works with GS to produce 19*Z*-geissoschizine shows how this protein also controls the stereochemistry of the product. We demonstrate that FoGS and GS interact to form a stable heterodimer, and show through mutagenesis that FoGS likely modulates the structure of the GS active site. Microscopy studies suggest that FoGS also changes the subcellular localization of GS. The discovery of FoGS will enable efforts to heterologously reconstitute commercially important geissoschizine-derived monoterpene indole alkaloids, and moreover, highlights how metabolic pathways utilize proteins that are not essential for production of the products, but nevertheless enhance the efficiency and specificity of the catalytic biosynthetic pathway components.

## Introduction

Monoterpene indole alkaloids are an important class of natural products produced by plant species in the order Gentianales, with over 3000 structurally distinct molecules reported (1). The medicinal value of monoterpene indole alkaloids is exemplified by compounds such as vinblastine and vincristine (anticancer drugs) (2, 3), ibogaine (antiaddictive activity) (4, 5), ajmalicine (antihypertensive activity) (6), strychnine (pesticide) (7), and quinine (antimalaria drug) (8). Given the structural complexity and pharmacological importance of these alkaloids, substantial efforts have been made to elucidate the pathways that encode the biosynthesis of these compounds. All monoterpene indole alkaloids are derived from strictosidine, an intermediate that is formed by enzyme-catalyzed condensation of tryptamine and the monoterpene secologanin (9). Strictosidine is degylcosylated by strictosidine β-glucosidase (SGD) to form a labile aglycone that can rearrange to form a variety of isomers (10, 11). These strictosidine aglycone isomers are reductively trapped by medium-chain dehydrogenase/reductases to generate the scaffolds for the major classes of monoterpene indole alkaloids (12–18) (Figure 1). A particularly important strictosidine aglycone isomer is dehydrogeissoschizine, which is reduced by geissoschizine synthase (GS) to form 19*E*-geissoschizine (12, 17), a precursor for strychnos, aspidosperma and iboga type alkaloids, which include many medicinally important compounds including vinblastine, strychnine and ibogaine.

**Figure 1:**
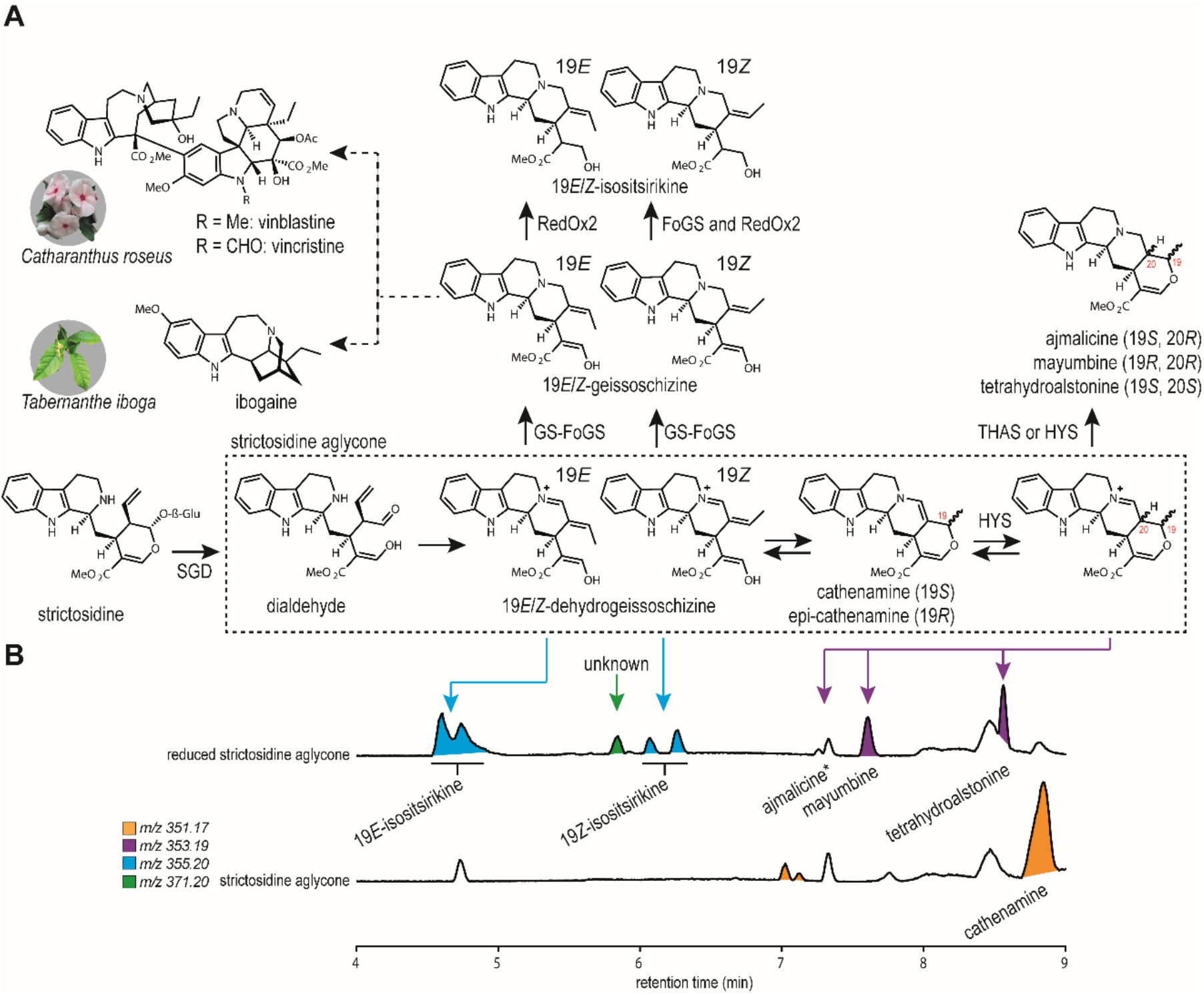
The strictosidine aglycone branch point. **A** The enzymes and intermediates of the strictosidine aglycone branchpoint in *C. roseus* and *T. iboga* pathways. Major strictosidine aglycone isomers are shown within the dotted line. Solid arrows represent single enzymatic/chemical conversions and dotted arrows represent multi-enzyme steps. SGD, strictosidine β-glucosidase; GS, geissoschizine synthase; FoGS, facilitator of geissoschizine synthase; HYS, heteroyohimbine synthase; THAS, tetrahydroalstonine synthase **B** LC-MS baseline peak chromatograms of strictosidine aglycone and the products of subsequent reduction by sodium borohydride. Peaks are highlighted based on *m/z* values. Ajmalicine is present at trace levels, but is obscured by a larger peak. Arrows are drawn to show the most likely strictosidine aglycone species that the subsequent reduction product corresponds to. Double peaks of 19*E*- and 19*Z*-isositsirikine correspond to 16*R* and 16*S* stereoisomers, previously described (23).

Herein we report the discovery of an auxiliary protein, *facilitator of geissoschizine synthase* (FoGS), that aids in the formation of geissoschizine through heterodimerization with GS. The term *auxiliary protein* is used since FoGS appears to enhance the efficiency and specificity of the reduction, but does not directly catalyze the reaction. We identified orthologues of FoGS from the geissoschizine producers *Catharanthus roseus* and *Tabernanthe iboga* and demonstrate activity for all orthologues in *Nicotiana benthamiana* and *in vitro*. Virus induced gene silencing (VIGS) demonstrate the clear involvement of FoGS in the geissoschizine-derived vinblastine biosynthetic pathway in *C. roseus*. FLIM-FRET and co-immunoprecipitation experiments showed protein-protein interactions between FoGS and GS, and suggest that the auxiliary activity is mediated by a heterodimerization of GS and FoGS. The discovery of FoGS will enable heterologous production of geissoschizine-derived monoterpene indole alkaloids (e.g. vinblastine, ibogaine). Moreover, this discovery highlights how metabolic pathways utilize proteins that are not essential, but nevertheless enhance the titers of certain biosynthetic intermediates. The availability of highly resolved datasets and high-throughput functional characterization screens will likely facilitate discovery of more examples of auxiliary proteins.

## Results

### A medium-chain dehydrogenase/reductase facilitates *Catharanthus roseus* geissoschizine synthase

In *C. roseus*, strictosidine aglycone isomers are reductively trapped by one of the medium-chain dehydrogenases/reductases GS, HYS, or THAS (12, 15, 14, 17). GS catalyzes conversion of strictosidine aglycone (reduction of dehydrogeissoschizine isomer) to 19*E-* geissoschizine, which is directed to strychnos/aspidosperma/iboga-type alkaloids. Alternatively, HYS and THAS reduce strictosidine aglycone (cathenamine isomer) to the heteroyohimbine-type alkaloids ajmalicine (HYS), tetrahydroalstonine (THAS or HYS) or mayumbine (HYS). Inspired by the chalcone isomerase-like proteins that enhance titers of flavonoid biosynthetic pathways (19– 21), we speculated that a non-reductive protein could assist the efficiency or specificity of the isomerization process involved in this step of the monoterpene indole alkaloid pathway.

Using a published *C. roseus* single-cell transcriptome dataset (22), we identified candidate genes based on co-clustering with SGD in epidermal cells (Figure S1). Candidates were screened in batches via agrobacterium-mediated expression in *N. benthamiana*. We tested all candidates in the presence of strictosidine and SGD to generate the strictosidine aglycone substrate. Additionally, we included the three major medium-chain dehydrogenases/reductases that act on strictosidine aglycone, GS1, HYS, or THAS1. This competitive assay would enable us to identify auxiliary proteins that would alter the subsequent alkaloid product profiles: ajmalicine, mayumbine, and tetrahydroalstonine for HYS; tetrahydroalstonine for THAS1; and 19*E*-geissoschizine for GS1. In *N. benthamiana*, the enol moiety of geissoschizine stereoisomers (19*E* or 19*Z*) is further reduced by an endogenous reductase or reductases to form 19*E*- and 19*Z*-isositsirikine while retaining the original 19*E*/*Z*-stereochemistry set by GS (23) (Figure S2).

These *N. benthamiana* assays established that ajmalicine is the major product of HYS, a stark difference to the even distribution of ajmalicine, mayumbine, and tetrahydroalstonine that is produced by HYS in *in vitro* assays (15). We additionally noted that in this competitive assay, GS1 is outcompeted by HYS, reflected by the higher levels of ajmalicine compared to the 19*E*-geissoschizine derived isositsirikine. Notably, one of the auxiliary protein candidates increased flux through GS1 at the expense of HYS, as evident by an increase in 19*E*-isositsirikine and decrease in ajmalicine (Figure 2A and S1). This candidate, which was annotated as a medium-chain dehydrogenase/reductase, was unable to reduce strictosidine aglycone when expressed alone with SGD in the presence of strictosidine. We thus named this candidate *facilitator of geissoschizine synthase* (FoGS). FoGS was then assayed with each medium-chain dehydrogenase/reductase individually. Assay of strictosidine, SGD, FoGS and either GS1, HYS, and THAS1 revealed no significant changes to the resulting product profiles (Figure S3). Since omission of THAS1 from competition assays had no effect on product profiles (Figure S3), we omitted THAS1 from subsequent experiments. This is congruent with single cell transcriptomics data that show THAS1 is not expressed in epidermal cells and is therefore spatially separated from SGD and GS1 (22, 24) (Figure S1).

**Figure 2:**
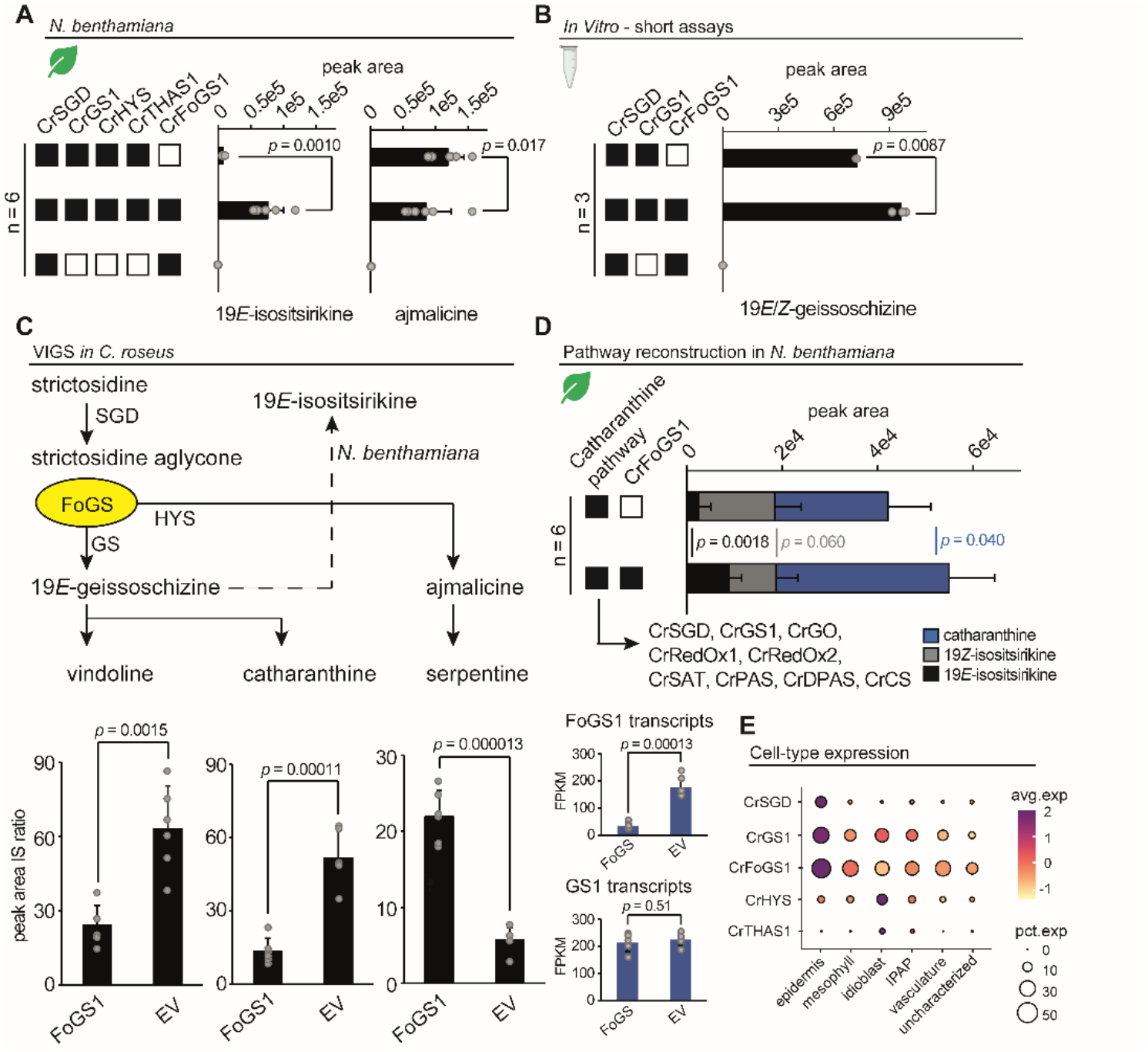
A medium-chain dehydrogenase/reductase facilitates the GS1 enzyme in *Catharanthus roseus*. For all bar graphs, values for individual replicates are shown as grey circles and averages as black bars. Error bars and *p*-values from student’s t-tests are shown. **A** Competitive SGD, GS1, HYS, THAS1 assays in *N. benthamiana* with and without FoGS1. Results for other candidates are shown in Figure S1. Each treatment is the average of 6 biological replicates. **B** *In vitro* characterization of GS1 and FoGS1. Combined peak areas of 19*E*- and 19*Z*-geissoschizine from three replicates are shown. **C** Virus induced gene silencing (VIGS) of FoGS1 in *C. roseus* leaves. A diagram highlighting the major enzymes, intermediates, and downstream products of the strictosidine aglycone branchpoint. Results from targeted metabolomics of vindoline, catharanthine, and serpentine are shown below the diagram alongside average FPKM values for FoGS1 and GS1 from 3 representative samples. Average peak areas were determined from 6 biological replicates. **D** Reconstruction of the *C. roseus* 8-step biosynthetic pathway from strictosidine to catharanthine in *N. benthamiana* leaves. Each treatment is the average of 6 biological replicates. Blue corresponds to catharanthine, grey to 19*Z*-isositsirikine, and black to 19*E*-isositsirikine. Full dataset showing individual replicate values are shown in Figure S4. **E** Single cell transcriptomics for selected *C. roseus* genes. Although HYS is predominantly expressed in idioblast cells, HYS is expressed in both epidermal and idioblast cells under certain conditions (24). Single cell transcriptomics are represented by a dotplot where circle size corresponds to percent expression and circle colour corresponds to average expression. SGD, strictosidine β-glucosidase; GS, geissoschizine synthase; FoGS1, facilitator of geissoschizine synthase; HYS, heteroyohimbine synthase; THAS, tetrahydroalstonine synthase; GO, geissoschizine oxidase; RedOx1, reductive oxidative enzyme 1; RedOx2, reductive oxidative enzyme 2; SAT, stemmadenine acetyl transferase; PAS, precondylocarpine acetate synthase; DPAS, dehydroprecondylocarpine synthase; CS, catharanthine synthase.

To validate whether FoGS plays a role in 19*E*-geissochizine synthesis *in planta*, we silenced FoGS in *C. roseus* leaves by virus induced gene silencing (VIGS). Silencing of FoGS resulted in an 80% decrease in transcript levels compared to empty vector controls without altering GS1 expression. This was accompanied by a decrease in the 19*E*-geissoschizine-derived metabolites catharanthine (iboga type), vindoline (aspidosperma type), and an increase in the ajmalicine-derived metabolite serpentine (heteroyohimbine type) (Figure 2C). Therefore, the VIGS data clearly support a role for FoGS in directing metabolic flux at the GS-HYS branch point that controls the switch between aspidosperma/iboga and heteryohimbine-type alkaloids. The FoGS VIGS chemotype closely resembles what is observed when GS1 is subjected to silencing, for which a decrease in 19*E*-geissoschizine-derived metabolites and an increase in ajmalicine is also reported (25). These results suggest that FoGS enhances 19*E*-geissoschizine production at the strictosidine aglycone branch point.

### *In vitro* characterization of *Cr*FoGS

To further support the role of FoGS, we expressed and purified SGD, GS1, and FoGS recombinantly for *in vitro* characterization. The *in vitro* enzyme assays were conducted by first deglycosylating strictosidine with SGD and allowing the aglycone to equilibrate before the addition of GS1 and FoGS. LC-MS analysis of equilibrated strictosidine aglycone revealed only one major peak with an *m/z* value of 351 (Figure S5). This product was purified by HPLC and NMR analysis indicated that this product was the strictosidine aglycone isomer cathenamine (Figure S6), consistent with previous reports (10, 11, 26). Quenching this aglycone reaction with sodium borohydride, and comparison of the resulting product mixture with authentic standards, demonstrated that cathenamine can be chemically reduced to 19*E*-, 19*Z*-isositsirikine, mayumbine, tetrahydroalstonine, and ajmalicine (Figure 1B). These results suggest that cathenamine is in equilibrium with the aglycone substrates (19*E*-, 19*Z*-dehydrogeissoschizine and tautomerized iminium forms of cathenamine and epi-cathenamine) that lead to the products of GS1 and HYS. Importantly, *in vitro* enzyme assays show that FoGS modestly increases GS1 catalyzed production of 19*E*-(major) and 19*Z*-geissoschizine (minor) (Figure 2B). Interestingly, FoGS also displayed modest 19*Z*-isositsirikine synthase activity as evidenced by the reduction of 19*Z*-geissoschizine in coupled assays. This was confirmed with assays that incubated FoGS with synthetic 19*E*- and 19*Z*-geissoschizine as substrates (Figure S7). In extended assays, FoGS reduced strictosidine aglycone to form two unknown products with *m/z* values of 353.19 and 355.20, though the low titers of these products prevented isolation and structural characterization. These products were not observed in *N. benthamiana* assays (Figure 2B). Therefore, FoGS retains the catalytic capacity for reduction, though this activity does not appear to be utilized in geissoschizine production.

### *Tabernanthe iboga* FoGS orthologues provide added *E/Z*-stereochemical control

We next asked whether FoGS orthologues are utilized in other geissoschizine producing species. *Tabernanthe iboga*, which produces the antiaddiction agent ibogaine and is phylogenetically related to *C. roseus*, was targeted for analysis. GS from *T. iboga* is a catalytically inefficient enzyme that produces 19*Z*-geissoschizine as a major product, even though the downstream enzyme, geissoschizine oxidase (GO), only turns over 19*E*-geissoschizine (23). In our hands, *Ti*GS showed trace 19*Z*-geissoschizine synthase activity in *N. benthamiana* and no detected activity *in vitro* (Figure 3AB). We suspected that a FoGS orthologue may be required for production of 19*E*-geissoschizine by *Ti*GS. *Ti*FoGS candidates were selected based on sequence similarity with *Cr*FoGS1 via BLAST analysis of a published *T. iboga* transcriptome (27). Candidates were initially screened via agrobacterium-mediated expression in *N. benthamiana* with *Ti*SGD and *Ti*GS and strictosidine as a starting substrate. Again, since *N. benthamiana* reduces *E/Z*-geissoschizine, levels of this reduced product, *E/Z*-isositsirikine, were measured in the assays. Three candidates, *Ti*FoGS1, *Ti*FoGS2, and *Ti*FoGS3, all substantially increased the levels of geissoschizine produced by *Ti*GS, with varying degrees of 19*E*/*Z* specificity (Figure 3A). Addition of *Ti*FoGS1 to the assay enabled formation of 19*E*-geissoschizine by *Ti*GS, similar to *Cr*FoGS, thereby explaining how 19*E*-geissoschizine, required for ibogaine biosynthesis, is produced in this plant. *Ti*FoGS3 allowed *Ti*GS to produce 19*Z*-geissoschizine, while addition of *Ti*FoGS2 resulted in a mixture of 19*E*- and 19*Z*-geissoschizine. When expressed alone with *Ti*SGD, *Ti*FoGS1, *Ti*FoGS2, and *Ti*FoGS3 all showed low levels of GS-activity, comparable to the level of activity exhibited by *Ti*GS alone (Figure S8). FoGS activities were reproduced *in vitro* with purified protein (Figure 3B). *Ti*FoGS orthologues also converted strictosidine aglycone to the same unknown product (*m/z* 353.19) observed with *Cr*FoGS1 (Figure S9). Only *Ti*FoGS3 could reduce 19*Z*-geissoschizine to 19*Z*-isositsirikine (Figure 3B). Using a deposited single-cell transcriptome from *T. iboga* leaves (see supplemental methods), we observed that *Ti*FoGS1 is most clearly clustered with *Ti*SGD and *Ti*GS (Figure 3C). We therefore propose that *Ti*FoGS1 is the major contributor to GS-catalyzed production of 19*E*-geissoschizine in *T. iboga*.

**Figure 3:**
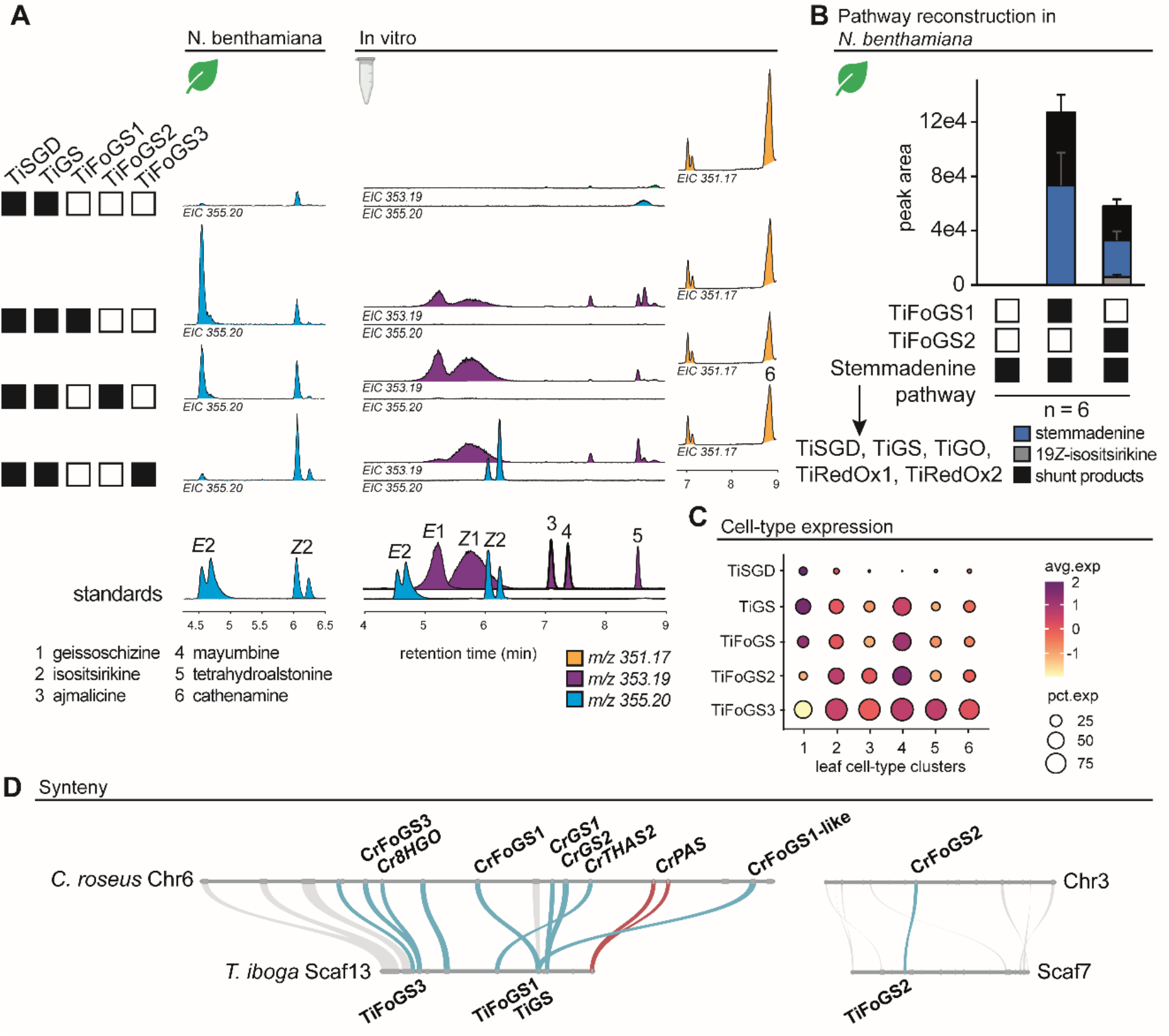
FoGS orthologues from *T. iboga*. **A** Characterization of *Ti*FoGS1, *Ti*FoGS2, and *Ti*FoGS3 with *Ti*SGD and TiGS in *N. benthamiana* and *in vitro*. Extracted ion chromatograms (EIC) are shown for *m/z* values of 353 (purple), 355 (cyan), and 351 (orange). Standards for 19*E*/*Z*-geissoschizine (1), 19*E*/*Z*-isositsirikine (2), ajmalicine (3), mayumbine (4), and tetrahydroalstonine (5) are shown below. **B** Reconstruction of the *T. iboga* 4-step *biosynthetic* pathway from strictosidine to stemmadenine in *N. benthamiana* leaves. Each treatment is the average of 6 biological replicates. Blue corresponds to stemmadenine, grey to 19*Z*-isositsirikine, and black to 19*E*-isositsirikine. Shunt products are formed endogenously in *N. benthamiana* from stemmadenine. Full dataset showing individual replicate values are shown in Figure S9. **C** Single cell transcriptomics for select *T. iboga* genes, represented by a dotplot where circle size corresponds to percent expression and circle colour corresponds to average expression. **D** Microsynteny comparison highlights *C. roseus* and *T. iboga* genomic regions containing FoGS genes. Blue lines indicate synteny between medium-chain dehydrogenases/reductases, red for flavin containing berberine bridge-like proteins (PAS), and grey for other. SGD, strictosidine β-glucosidase; GS, geissoschizine synthase; FoGS, facilitator of geissoschizine synthase; THAS, tetrahydroalstonine synthase; GO, geissoschizine oxidase; RedOx1, reductive oxidative enzyme 1; RedOx2, reductive oxidative enzyme 2; PAS, precondylocarpine acetate synthase.

### FoGS is located in a biosynthetic gene cluster with GS

Synteny analysis using published *C. roseus* (22) and *T. iboga* (not yet released) genomes revealed that *Cr*FoGS, *Ti*FoGS1 and *Ti*FoGS3 are located in a previously noted genomic region containing GS1, GS2, THAS1, as well as two additional monoterpene indole alkaloid genes (8HGO and PAS) (28) (Figure 3D). *Ti*FoGS1 and *Ti*FoGS3 are syntenic to *Cr*FoGS and *Cr*8HGO, respectively, alongside two uncharacterized medium-chain dehydrogenases/reductases. *Ti*FoGS2 is located outside the gene cluster and is syntenic to an uncharacterized *C roseus* medium-chain dehydrogenase/reductase. Single-cell transcriptomics of the newly identified *C. roseus* FoGS syntenologues reveal that only *Cr*FoGS1 is co-expressed in epidermal cells with *Cr*GS1. FoGS-activity was detected for *Cr*FoGS2 and *Cr*FoGS3 but not for *Cr*8HGO when assayed in in *N. benthamiana* (Figure S11). However, *Cr*FoGS2 and *Cr*FoGS3 did not provide the observed 19*E*/*Z*-stereochemical control exhibited by the *T. iboga* counterparts.

### Protein-protein interactions between *C. roseus* FoGS1 and GS1

We hypothesized that protein-protein interactions between FoGS1 and GS1 contribute to the observed auxiliary role of FoGS1. We first assessed the subcellular localization of the *C. roseus* strictosidine aglycone branch point enzymes, SGD, GS1, FoGS1, and HYS, by transiently expressing GFP/RFP fusion constructs in *N. benthamiana* leaves (Figures S12-15). *Cr*SGD and *Cr*HYS demonstrated similar subcellular localization patterns as previously reported (15, 26, 29). *Cr*SGD formed dense *fusiform* nuclear structures that were disrupted by C-terminal tagging (Figure S13), and *Cr*HYS displayed nucleocytosolic localization. *Cr*GS1 showed nucleocytosolic localization with a strong nuclear signal, while *Cr*FoGS1 displayed nucleocytosolic localization with a weak nuclear signal. Unexpectedly, when co-expressed with *Cr*FoGS1, the subcellular localization of *Cr*GS1 was modulated, appearing predominantly excluded from the nucleus, thereby matching the localization of *Cr*FoGS (Figure S14). *T. iboga* SGD, GS, and FoGS1 displayed near identical patterns of subcellular localization as the *C. roseus* orthologues when expressed in *N. benthamiana* (Figure S16). Importantly, we observed the same exclusion of GS from the nucleus when co-expressed with FoGS1 (Figure S17 and S18).

The shift in GS subcellular localization to align with the localization pattern of FoGS1 suggests that these two proteins interact. We used fluorescence lifetime imaging microscopy (FLIM) to detect Förster resonance energy transfer (FRET) between GFP and RFP protein fusions transiently expressed in *N. benthamiana* leaves. In pairwise FLIM-FRET assays between *C. roseus* GS1, FoGS1, HYS, and SGD, FRET was only observed for *Cr*GS-FoGS1 (4.8-5.5% FRET) and *Cr*GS-HYS (2.7-4.5% FRET) (Figure 4A and S20). We also note the unusual behaviour of C-terminally tagged SGD, for which FRET signal was measured in all assays including a free-GFP negative control (Figure S20). To validate the results of the FRET experiments, we performed co-immunoprecipitation from *N. benthamiana* leaves transiently expressing *C. roseus* SGD, GS1, FoGS1, and HYS using GFP-*Cr*GS1 as bait. Co-immunoprecipitated proteins were analyzed by SDS-PAGE and proteomic analysis. Proteomics showed that only *Cr*FoGS1 was strongly enriched with GFP-*Cr*GS1 in co-immunoprecipitation assays, while *Cr*HYS showed modest enrichment, and *Cr*SGD was not enriched at all (Figure 4B). Congruently, co-immunoprecipitation experiments using GFP-*Cr*SGD as bait demonstrated a lack of enrichment for *Cr*GS1, *Cr*FoGS1, and *Cr*HYS (Figure S21). We also observed the appearance of a strong protein band in the SDS-PAGE of GFP-*Cr*GS1 co-precipitated proteins. This band had the molecular weight of a medium-chain dehydrogenase/reductase and likely corresponds to *Cr*FoGS1 based on the proteomics data (Figure S21). Collectively, these results suggest that *Cr*FoGS1 and *Cr*GS1 form a stable complex.

**Figure 4:**
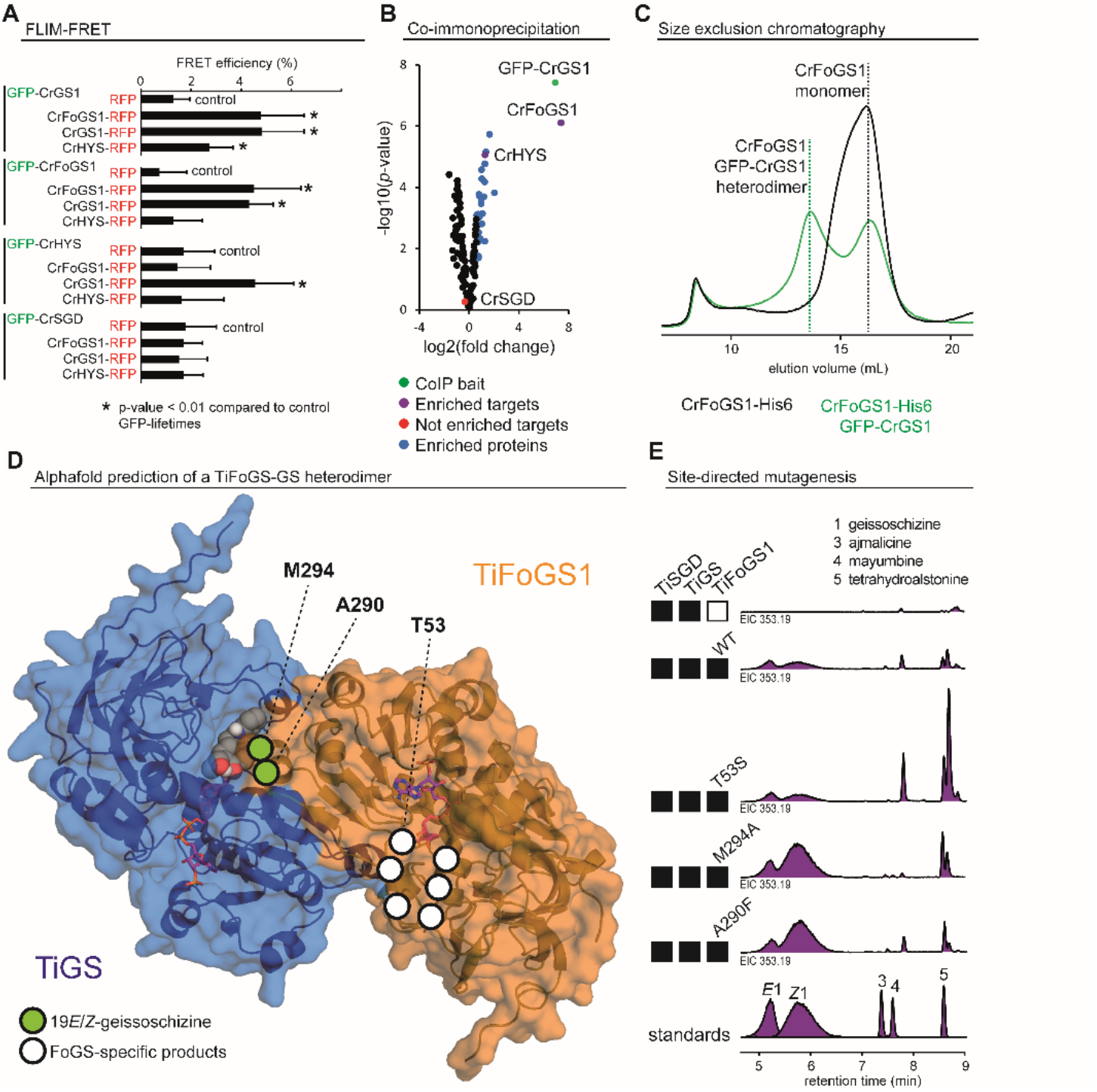
Protein-protein interactions between FoGS1 and GS. **A** Pairwise FLIM-FRET assays between *Cr*SGD, *Cr*GS1, *Cr*FoGS1, and *Cr*HYS expressed in *N. benthamiana* leaves. FRET efficiencies are shown as an average of 6 measurements from 3 biological replicates. Errors bars are shown. Treatments which had significantly different GFP fluorescence lifetimes than free-RFP negative controls (*p*-values from student’s t-tests < 0.01) are marked with an asterisk. **B** Volcano plot showing the enrichment of proteins between GFP-*Cr*GS1 Co-IPs and freeGFP negative controls. The results were determined from three biological replicates. GFP-*Cr*GS1 (bait) is shown in green, enriched proteins in blue, enriched proteins of interest in purple, and non-enriched proteins of interest in red. **C** Size exclusion chromatography of the purified *C. roseus* FoGS1-GS1 complex. Chromatogram for His^6^-*Cr*FoGS1 expressed alone is shown in black and His^6^-*Cr*FoGS1 co-expressed with GFP-*Cr*GS1 is shown in green. SDS-PAGE and Western blots of collected fraction and the calibration curve used to calculate molecular weight are shown in Figure S22. **D** Alphafold3 prediction (ipTM = 0.94; pTM = 0.93) of the *T. iboga* FoGS1-GS heterodimer. *Ti*GS is shown in blue, *Ti*FoGS1 in orange, NADPH in magenta, and docked 19*E*-dehydrogeissoschizine as grey spheres. The location of amino acids from site-directed mutagenesis of *Ti*FoGS1 and the corresponding effect of geissoschizine formation (green) and FoGS1-specific product formation (white) are mapped onto the model. **E** Site directed mutagenesis of *Ti*FoGS1. *In vitro* GS-coupled assays of *Ti*FoGS1 variants. Extracted ion chromatograms (EIC) are shown for *m/z 355*.*20* in purple. Standards for 19*E*/*Z*-geissoschizine (1), ajmalicine (3), mayumbine (4), and tetrahydroalstonine (5) are shown below. Full results from site-directed mutagenesis are shown in Figure S23. SGD, strictosidine β-glucosidase; GS, geissoschizine synthase; FoGS, facilitator of geissoschizine synthase; HYS, heteroyohimbine synthase.

### Heterodimerization of FoGS and GS

To investigate the molecular arrangement of the FoGS1-GS interaction, we purified the complex from *N. benthamiana* leaves co-expressing His^6^-*Cr*FoGS1 and GFP-*Cr*GS1. His^6^-*Cr*FoGS1 and interacting GFP-*Cr*GS1 were first captured using Ni-NTA affinity chromatography, then subsequently separated by molecular weight using size exclusion chromatography. When His^6^-*Cr*FoGS1 was expressed in the absence of GFP-*Cr*GS1, a single peak was observed during size exclusion chromatography with an elution volume corresponding to monomeric *Cr*FoGS1 (Figure 4C). In contrast, when His^6^-*Cr*FoGS1 was co-expressed with GFP-*Cr*GS1, an additional higher molecular weight peak was observed (Figure 4C). SDS-PAGE and Western blots confirmed that the additional peak contains both *Cr*FoGS1 and GFP-*Cr*GS1 (Figure S22). Importantly, the elution volume of this peak corresponds to that of a GFP-*Cr*GS1-*Cr*FoGS1 heterodimer. We observed the same peak distribution during size exclusion chromatography of purified *T. iboga* His^6^-FoGS1 and YFP-GS, demonstrating that FoGS1-GS heterodimerization is conserved in *C. roseus* and *T. iboga* (Figure S22). Modeling with Alphafold3 suggests that the FoGS1-GS heterodimer possesses the same quaternary structure as the empirically determined *Cr*GS1 homodimer (Figure 4D) (30). This mode of dimerization is also conserved in other medium-chain dehydrogenases/reductases, including examples from the monoterpene indole alkaloid pathway ((15, 30, 31), reviewed in (32)). The resulting quaternary organization produces two active sites that are predominantly located within each protomer but have a distinct contribution of a short α-helix from the neighbouring protomer. In this manner, the heterodimerization of FoGS1 and GS would result in two novel active sites: a GS active site with FoGS1 contribution, and a FoGS1 active site with GS contribution.

To test the predicted model, we made single amino acid substitutions to the canonical medium-chain dehydrogenase/reductase active site of FoGS1 and tested the effect in combination with wild-type GS *in vitro. T. iboga* enzymes were used because of the clear qualitative effect of *Ti*FoGS1 on *Ti*GS-activity. Active site residues were selected based on an Alphafold3 model of a *Ti*FoGS1 homodimer (Figure S23), with the intent to disrupt potential catalytic residues and change 19*E*/*Z* specificity. Only two positions (M294 and A290) resulted in a change in the relative levels of 19*E*- and 19*Z*-geissoschizine formation. M294A, M294Q, and A290F *Ti*FoGS1 variants all increased the levels of 19*Z*-geissoschizine and to a lesser extent 19*E*-geissoschizine (Figure 4E and S23). The T53S *Ti*FoGS1 mutant increased the levels of the uncharacterized product (*m/z* 353.19) that is produced when wild type *Ti*FoGS1 is incubated with strictosidine aglycone in the absence of *Ti*GS (Figure 4E). Mapping the tested *Ti*FoGS1 active site residues on the *Ti*FoGS1-*Ti*GS heterodimer model revealed that M294 and A290 are the only tested residues that contribute to the *Ti*GS active site (Figure 4D). Docking 19*E*-dehydrogeissoschizine into the *Ti*FoGS1-*Ti*GS heterodimer model highlights the contributions these residues make to the *Ti*GS active site, particularly M294, which likely interacts with the indole moiety of dehydrogeissoschizine (Figure S24). Collectively, these findings support a model in which the GS active site is modulated by heterodimerization with the auxiliary protein FoGS.

### Pathway reconstruction in *Nicotiana benthamiana*

We next tested whether FoGS could be leveraged to improve the heterologous pathway reconstruction in *N. benthamiana* leaves. To do this, we co-expressed the genes from *C. roseus* responsible for converting strictosidine to catharanthine in *N. benthamiana* with and without *Cr*FoGS1. The addition of *Cr*FoGS1 resulted in a modest increase of catharanthine and 19*E*-isositsirikine peak areas (Figure 2D). Additionally, the same experiment was conducted for the *T. iboga* genes responsible for converting strictosidine to stemmadenine using *Ti*FoGS1 and *Ti*FoGS2. In this case, *Ti*FoGS1 resulted in the greatest increase in stemmadenine peak area compared to *Ti*FoGS2, for which 19*Z*-isositirikine was also observed (Figure 3B). This result is consistent with the 19*E*- and 19*Z*-geissoschizine stereospecificity imparted to *Ti*GS by *Ti*FoGS2. Collectively, these results reinforce the biosynthetic role of FoGS and demonstrate its potential to improve, albeit modestly, titers of valuable monoterpene indole alkaloids in heterologous expression systems.

## Discussion

Here we report the discovery of an auxiliary protein, FoGS, that enhances the efficiency and specificity of the central monoterpene indole alkaloid biosynthetic gene GS. Orthologues of FoGS were identified in two related plants, *C. roseus* and *T. iboga*, both of which produce geissoschizine. Inclusion of the *Cr*FoGS1 protein only modestly impacted the yields of the *C. roseus* GS1 orthologue in a stand-alone assay. However, when *Cr*FoGS1 was assayed in a competition assay in which strictosidine aglycone was incubated with multiple enzymes that competes for the strictosidine aglycone substrate (*Cr*HYS and *Cr*THAS1), higher relative levels of the *Cr*GS1 product were observed. Moreover, silencing of *Cr*FoGS1 in *C. roseus* leaves showed that absence of this protein significantly lowers the levels of downstream geissoschizine derived metabolites *in planta*. The *T. iboga* GS orthologue absolutely requires FoGS1 to catalyze the formation of 19*E*-geissoschizine. Interestingly, we identified an additional *T. iboga* orthologue, *Ti*FoGS3, that imparts 19*Z*-stereochemical control over *Ti*GS. A biological role for *Ti*FoGS3 in geissoschizine biosynthesis is unlikely since *Ti*GS and *Ti*FoGS3 are not co-expressed in the same cell-types. However, the stereochemical control imparted to *Ti*GS by *Ti*FoGS1 and *Ti*FoGS3 provides an intriguing opportunity to access 19*E*- and 19*Z*-geissoschizine for biotechnological applications.

FLIM-FRET and co-immunoprecipitation experiments convincingly showed that *Cr*FoGS1 and *Cr*GS1 interact. Additionally, *C. roseus* and *T. iboga* FoGS1-GS heterodimers could be purified from *N. benthamiana* leaves. The quaternary organization of FoGS1 and GS in the heterodimeric structure that is predicted by Alphafold3 suggests that FoGS1 comprises part of the active site of GS. This prediction is supported by site-directed mutagenesis in which only residues at the predicted FoGS1-GS interface influenced geissoschizine formation. Collectively, our results suggest that native geissoschizine synthase activity in *C. roseus* and *T. iboga* is provided by a heterodimer of FoGS1 and GS, and not GS alone.

The discovery of the auxiliary protein, FoGS, highlights the potential for missing proteins in monoterpene indole alkaloid biosynthesis. As the biosynthetic routes to many of these valuable compounds are elucidated (10, 12, 18, 25, 34–36), additional factors that contribute to alkaloid biosynthesis may be discovered. Indeed, numerous auxiliary proteins have been reported to contribute to the biosynthesis of other plant natural products. In flavonoid biosynthesis, chalcone isomerase-like (CHIL) rectifies derailment caused by lactonization during the chalcone synthase (CHS) catalyzed formation of naringenin chalcone (19–21). *Facilitator of taxane oxidation* (FoTO1), was shown to eliminate shunt products not conductive to downstream biosynthesis during the oxidation of taxadiene from taxol biosynthesis in yew (37). Both CHIL and FoTO1 were shown to interact with the corresponding partner enzyme (20, 21, 37). Auxiliary proteins may also act by binding metabolites, as exemplified by the alkaloid-binding major latex proteins (MLPs) from opium poppy (38) and the flavonoid-binding pathogenesis related 10 proteins (PR10s; analogous to MLP) from strawberry (39). Recently, a PR10 from *Marchantia polymorpha* was shown to protect naringenin chalcone from spontaneous cyclization in flavonoid biosynthesis (40). Non-catalytic proteins in lignin (41) and steroidal alkaloid biosynthesis (42) are hypothesized to enhance product titers by acting as a scaffold for the biosynthetic enzymes. Core eudicots synthesize indole by utilizing an inactive paralogue of the tryptophan synthase β subunit (TSB-like), which in turn disrupts the canonical tryptophan synthase multimer (43). Additionally, in *C. roseus*, an alternate and inactive splice variant of SGD was shown to inhibit regular SGD function (26).

The discovery of the auxiliary protein FoGS has important implications for improving the heterologous production of high value geissoschizine derived alkaloids such as vinblastine (*C. roseus*) and ibogaine (*T. iboga*). Moreover, this discovery highlights the involvement of proteins in metabolic pathways that, while not essential for catalysis, nevertheless increase the efficiency or specificity of a given metabolic step. Furthermore, the use of FoGS to control the stereochemistry of the 19-position of geissoschizine can be applied to leverage known monoterpene indole alkaloid biosynthetic pathways for biotechnological applications.

## Materials and Methods

Additional materials and methods are described in the supporting information.

### Candidate selection

Candidates were selected using a previously published *C. roseus* single cell RNA-Seq leaf dataset (22) that was re-processed (alignment, clustering, and cell annotation) as previously reported. Transcripts annotated as *major latex proteins, pathogenesis related 10 proteins, alpha/beta-hydrolases*, or *medium-chain dehydrogenase/reductases* (including *cinnamyl-aldehyde*/*alcohol dehydrogenases*) were targeted. Major latex proteins and pathogenesis related 10 proteins were selected based on reported alkaloid binding activities (38, 39), and medium-chain dehydrogenase/reductases were selected because of the roles of GS, HYS, THAS, DCS, and YOS in reducing strictosidine aglycone (12–16, 18). Alpha/beta-hydrolases were selected based on reported cyclase activities in MIA biosynthesis (35). An initial list of 222 candidates was filtered based on Pearson correlation with SGD cell type expression, and the level of expression in epidermal clusters. Only candidates with no reported MIA biosynthetic roles that showed a Pearson correlation with SGD expression greater than 0.5 and exhibited expression in the epidermal cell cluster greater than 1 (average normalized counts per 10,000) were selected for screening. This yielded 6 major latex proteins/pathogenesis related 10 proteins, 3 alpha/beta-hydrolases, and 3 medium-chain dehydrogenases/reductases (Figure S1).

### Gene screening and pathway reconstruction in *N. benthamiana* leaves

Two-days post agrobacterium infiltration, *N. benthamiana* leaves transiently expressing the genes pertaining to the given experiment were infiltrated with 150 μM substrate in 50 mM HEPES pH 7.5 into the abaxial side of leaves. Leaf discs were excised three-days post substrate infiltration and metabolites were extracted in 400 μL methanol with gentle shaking. Samples were filtered through 0.45 μm low-binding hydrophilic polytetrafluoroethylene spin-filter plates (*Millipore*) and directly analyzed by liquid-chromatography – mass spectrometry (LC-MS). Analytes were verified with authentic standards based on retention time and MS/MS data (Figure S25). *N. benthamiana* assays were performed with 4-6 biological replicates, treating individual leaves from different plants as biological replicates.

### *In vitro* enzyme assays

Coupled SGD-GS enzyme assays were performed stepwise by first deglycosylating strictosidine with SGD then adding GS with and without FoGS. In the first step, 200 μM of strictosidine was degylcosylated by 20 nM SGD in 50 mM HEPES pH 7.5 and given 2 hours to equilibrate at 30 °C. We identified cathenamine as the major product of this reaction (Figure S5 and S6). Cathenamine reaction mixtures were then used as substrate for 50 μL enzyme assays of 0.25 μM GS, 0.1 μM FoGS, 1 mM NADPH, 0.1 mg/mL BSA, and 50 mM HEPES pH 7.5 at a final cathenamine concentration of 100 μM (assuming 100% conversion from strictosidine). Mixtures of GS and FoGS were pre-equilibrated for 30 min at 30 °C before the addition of cathenamine. *C. roseus* assays were run for 10 min while *T. iboga* assays were run for 1 hour, both at 30 °C. Extended assays of *C. roseus* FoGS1 in the absence of GS were run for 1 hour. *C. roseus* FoGS1 19*E*- and 19*Z*-geissoschizine assays were run for 1 hour using the same conditions with 50 μM geissoschizine. All assays were quenched with 4-volumes of 100 % methanol, vortexed, then centrifuged for 30 min at 4 °C to pellet proteins. Samples were filtered through 0.45 μm low-binding hydrophilic polytetrafluoroethylene spin-filter plates (*Millipore*) and analyzed using the same LC-MS method as for *N. benthamiana* assays. *In vitro* enzyme assays were performed in triplicate and analytes were verified with authentic standards based on retention time and MS/MS data (Figure S25).

### Fluorescence Lifetime Imaging Microscopy – Forster Resonance Energy Transfer (FLIM-FRET)

FLIM-FRET was performed as previously described (44). Briefly, eGFP tagged (N-terminal) and mRFP1 tagged (C-terminal) genes of interest were overexpressed in *N. benthamiana* leaves. Leaf discs were excised 3-days post agrobacterium infiltration and mounted in water. FLIM imaging was performed using Time-Correlated Single-Photon Counting (TCSPC) as implemented on a Stellaris 8 FALCON STED (*Leica*) equipped with a HC PL APO 40x/1.25 glycerol immersion objective (Leica). GFP was excited using a White Light Laser (WLL) (Leica) at a wavelength of 489 nm with 80 MHz pulse frequency. GFP emission between 495 and 583 nm was recorded using a HyD-X detector until reaching a photon/pixel count of 1000. An additional channel was used to image RFP fluorescence using an excitation wavelength of 590 nm and emissions between 597 and 640 nm were recorded using a HyD-S detector. GFP fluorescence lifetimes were determined from phasor plots (45), as implemented in Leica Application Suite X v.4.8.1. FRET efficiencies were calculated with equation 1 (46) from 6 biological replicates. *τ*_*unquenched*_ corresponds to the fluorescence lifetime of eGFP alone, and *τ*_*quenched*_ corresponds to the fluorescence lifetime of eGFP in the presence of the acceptor fluorophore mRFP1.

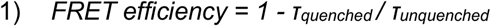

### Co-immunoprecipitation

Co-immunoprecipitation of GFP tagged (N-terminal) *Cr*GS1 and *Cr*SGD with the strictosidine aglycone branch point enzymes (*Cr*SGD, *Cr*GS1, *Cr*FoGS1, and *Cr*HYS) were performed using a µMACS™ GFP isolation kit (Miltenyi Biotech) following the manufacturer’s instructions. Briefly, GFP tagged (N-terminal) *Cr*GS1, *Cr*SGD, and untagged GFP were overexpressed in *N. benthamiana* leaves with *Cr*SGD, *Cr*GS1, *Cr*FoGS1, and *Cr*HYS. Leaves were harvested 3-days post agrobacterium infiltration, flash frozen in liquid nitrogen, and grounded by mortar and pestle. Proteins were extracted from 200 mg of leaf tissue in 0.5 mL of lysis buffer (150 mM NaCl, 1 % Ecosurf, 50 mM Tris-Cl pH 8) with an EDTA-free protease inhibitor cocktail for 1 hour at 4 °C with gentle shaking. Cell debris was removed by centrifugation at 14,000 rpm. 50 µL of anti-GFP magnetic beads were added to the supernatant and incubated for 30 min at 4 °C with gentle shaking. Samples were loaded onto µ Columns (*Miltenyi Biotech*) equilibrated in lysis buffer. Samples were washed four times with 200 µL and wash buffer 1 and once with 100 µL wash buffer 2. Proteins were incubated for 5 min with 20 µL of hot SDS-elution buffer then eluted with 50 µL of hot SDS-elution buffer. Samples were subjected to SDS-PAGE and GFP or GFP-tagged proteins were detected by Western Blotting using a GFP polyclonal antibody-horseradish peroxidase conjugate (Invitrogen) at a 1:1000 dilution followed by detection using Clarity Western ECL Substrate (*Bio-Rad*). Proteomics analysis was performed at the Proteomics core Facility, EMBL (Heidelberg, Germany). Co-immunoprecipitations were done in triplicate.

### Purification of the GS-FoGS1 complex

Open reading frames from *Cr*FoGS1 and *Ti*FoGS1 pOPINF constructs, including N-terminal His^6^ tags, were PCR amplified and assembled into pCambia vectors by In-Fusion. His^6^-FoGS was co-expressed with GFP/YFP tagged (N-terminal) GS in *N. benthamiana* leaves. For each treatment, 3 leaves from 10 plants were infiltrated. 3-days post agrobacterium infiltration, leaf tissue was flash frozen in liquid nitrogen and grounded by mortal and pestle. Pulverized plant tissue was incubated with 40 mL of buffer A1 supplemented with an EDTA-free protease inhibitor cocktail for 2 hours at 4 °C with gentle shaking. Cell debris was removed by centrifugation at 35,000 x g at 4 °C for 30 min, then filtered using Ministart^®^ NML Plus high-capacity syringe filters (*Sartorius*). Recombinantly expressed proteins were purified using an AKTA pure FPLC (*Cytiva*). Cleared lysate was loaded onto a His-Trap HP 5 mM column (*Cytiva*) at 2 mL/min equilibrated in buffer A1, and washed with 20 column volumes of buffer A1. Isocratic elution of his^6^-tagged proteins was performed with 100 % buffer B1 and 1.5 mL fraction were collected. Fractions corresponding to a single peak detected by UV at 280 nm were pooled. Size exclusion chromatography was then used to determine the molecular weight of proteins in the pooled fractions. Protein (400 µL) was injected onto a Superdex 200 10/300 GL size exclusion column (*Cytiva*) equilibrated in buffer A4 with a constant flow of 0.5 mL/min collecting 1 mL fractions. Eluting proteins were detected by UV at 280 nm. Fractions from size exclusion chromatography were analyzed by SDS-PAGE and GFP-tagged *Cr*GS1 detected by Western Blotting, as for co-immunoprecipitations. Molecular weights were calculated using a 4-point standard curve consisting of Catalase (240kDa), monomeric and dimer BSA (132 and 66 kDa), and Carbonic Anhydrase (30kDa). All elution volumes used to calculate molecular weights were determined from triplicate injections.

## Acknowledgments

We acknowledge Dr. Gyumin Kang for providing valuable chemicals and authentic standards. We also acknowledge Dr. Allwin McDonald, Sönke Beewen and Dr. Ling Chuang for advice and thoughtful discussion relevant to this work and Anja David for cloning assistance. We acknowledge funding from the Max Planck Society, DfG Leibniz Award (505457618).

