## Supplementary material for "An auxiliary protein tunes reductase activity in alkaloid biosynthesis": SI

**Contents:**

Supplemental Materials and Methods
Figures S1 to S26
Tables S1 to S5
SI References

**Supplemental Materials and Methods**

**Plants and plant growth**

*N. benthamiana* plants used for transient gene expression were cultivated in a greenhouse at 22 °C and 55% humidity with a 16-hour light and 8-hour dark photoperiod. Plants were transferred to growth chambers at 3–4-weeks of age with the same conditions prior to agrobacterium infiltration. *Catharanthus roseus* cultivar 'Atlantis Burgundy Halo' were cultivated in growth chambers at 21-28 °C and 60% humidity with a 16-hour light and 8-hour dark photoperiod. Plants were watered periodically as needed. *T. iboga* was obtained from the global ibogaine therapy alliance in May 2014, and since maintained as previously described (1).

**Chemicals and standards**

All chemicals used in this study were purchased as molecular biology grade or higher from commercial vendors (*Sigma Aldrich*, *Thermo Fischer*, etc.) unless stated otherwise. Ajmalicine and tetrahydroalstonine were purchased from *Sigma Aldrich*, catharanthine from *Abcam*, and ajmaline from *Extrasynthese*. 19*E*-geissoschizine, 19*E*-isositsirikine, and stemmadenine were from by Dr. Mohamed Omar Kamileen, as reported (2). 19*Z*-geissoschizine, 19*Z*-isositsirikine, and strictosidine were from Dr. Gyumin Kang, as reported (2).

**Single-cell transcriptome data processing and visualization**

For re-processing of previously published *C. roseus* single cell RNA-Seq leaf dataset, alignment, clustering, and cell annotations were performed as previously described (3). For processing *T. iboga* single cell RNA-Seq leaf dataset generated from a recent deposited dataset (SRR35790638, SRR35790637), sequencing reads were aligned to the *T. iboga* v1 genome (not yet released). Briefly, barcoded fastq files were generated from sequencing files by extracting cell barcodes based on the pipseeker pipeline (v.3.1.3, Illumina) and concatenated. The resulting files served as input to map reads to the genome via starsolo (v.2.7.10a), with the parameters: --soloCellFilter EmptyDrops_CR, --soloFeatures GeneFull, --soloMultiMappers EM (4). Raw and filtered matrices were used in SoupX (v.1.6.2) to assess the level of ambient RNA and remove them in dataset (5). The generated output file was used in Seurat (v.5.0.1) for downstream analyses (6). Cells were excluded from the analysis when less than 300 genes or more than 5,000 genes were detected in root dataset, and less than 500 genes and more than 5,000 genes were detected for leaf. Counts were log-normalized. 3,000 variable genes were selected and used for finding anchors based on CCA to integrate replicates. Data were scaled and PCA was performed using the variable features. UMAP was calculated using the first 30 principal components with default parameters. Expression level of each gene at cell type level were visualized as a dotplot, implementing DotPlot function in Seurat. Average expression in cell type was extracted from RNA slot of Seurat object and plotted in color-scale. Size of circles represent the percentage of cells expressing corresponding gene in cell type. FPKM values of each gene from *C. roseus* mRNA-seq datasets (SRR1271857, SRR1271858, SRR1271859) were extracted from Supplementary Table 4 of a previous study (3).

**Molecular biology**

Total RNA from *C. roseus* and *T. iboga* young leaves was extracted using a RNeasy Mini Kit (*Qiagen*) according to the manufacturer's instructions with an added on-column DNAse digestion. RNA quantification and purity were determined using a Implen NanoPhotometer N60. cDNA was prepared using Superscript IV VILO master mix. Full-length *C. roseus* SGD, GS1, HYS, THAS1, gene candidates and *T. iboga* gene candidates were amplified from young leaf cDNA (primers in Table S1). Gel purified PCR products were cloned into a modified 3omega1 (*BsaI*) (7), pOPINF (*HindIII* and *KpnI*), pCambia eGFP (*KpnI* and *PstI*), or pCambia mRFP1 (*KpnI* and *SalI*) vector employing the listed restriction sites for In-Fusion assembly (*Takara*). pCambia eGFP and mRFP1 vectors were constructed from pCambia split-luciferase vectors (8) by replacing the split-luciferase inserts with synthetic gene fragments of eGFP1 (GeneBank AAB02572) and mRFP1 (GeneBank CAH64892). *Ti*FoGS1 mutants were generated using overlap extension PCR and assembled using In-Fusion (Primers in Table S2). Assemblies were transformed into *Escherichia coli* Top10 competent cells and purified plasmids from single colonies were sequence-verified using Sanger sequencing. All agar and media were supplemented with 100 µg/mL spectinomycin (3omega1), 100 µg/mL kanamycin (pCambia) or 100 µg/mL carbenicillin (pOPINF).

Constructs for *Ti*SGD, *Ti*GS, *Ti*GO, *Ti*RedOx1, and *Ti*RedOx2 were provided by Dr. Mohamed Omar Kamileen, as reported (2). 3alpha1 *N. benthamiana* expression constructs for *Cr*GO, *Cr*RedOx1, *Cr*RedOx2, *Cr*SAT, *Cr*PAS, *Cr*DPAS, and *Cr*CS were provided by Dr. Dagny Grzech, as reported (9). Sequences for *C. roseus* and *T. iboga* FoGS1, FoGS2, and FoGS3 are listed in Table S3 and S4.

**Agrobacterium-mediated transient gene expression in *N. benthamiana***

3omega1 and pCambia constructs were transformed into electrocompetent *Agrobacterium tumefaciens* GV3101 (*Goldbio*) via electroporation. Cells were recovered for 3 hours at 28 °C before plating on LB supplemented with 20 µg/mL rifampicin, 50 µg/mL gentamycin and 50 µg/mL spectinomycin (3omega1) or 50 µg/mL kanamycin (pCambia). Single colonies were used to inoculate 10 mL liquid LB cultures, supplemented with appropriate antibiotic markers, and grown overnight at 28 °C with 200 rpm shaking. Cells were harvested by centrifugation at 4000 x g for 10 min and pellets were resuspended in infiltration buffer (10 mM MES, 10 mM $MgCl_2$, pH 5.6, 200 µM acetosyringone). Resuspended cells were incubated for 3 hours in the dark with gentle shaking. Equiproportional culture mixtures were prepared ensuring not to exceed a combined $OD_{600}$ of 0.8. Mixes were then infiltrated into the abaxial side of a leaf of a 3–4-week-old *N. benthamiana* plant.

**Virus induced gene silencing in *C. roseus***

*Cr*FoGS1 VIGS was performed as previously described (3). Briefly, a 300 bp fragment from the coding region of the *C. roseus* FoGS1 gene (CRO-06G024580) was amplified from the 3omega1-*Cr*FoGS1 construct and assembled into pTRV2-MgChl (10) using *Bam*HI and *Xho*I restriction sites for In-Fusion assembly. Primers are listed in Table S5. To avoid off-target gene silencing, the SNG VIGS tool (https://vigs.solgenomics.net/, https://doi.org/10.1016/j.molp.2014.11.024) was used to select the 300 bp VIGS target region. Electrocompetent *Agrobacterium tumefaciens* GV3101 (*Goldbio*) cells were transformed with pTRV2-MgChl and pTRV2-FoGS1-MgChl constructs. Single colonies were used to inoculated 3 mL LB media supplemented with 20 µg/mL rifampicin, 50 µg/mL gentamycin and 50 µg/mL kanamycin and grown overnight at 28 °C with 200 rpm shaking. Cells were harvested by centrifugation at 4000 x *g* for 10 min and pellets were resuspended in infiltration buffer (10 mM MES, 10 mM $MgCl_2$, and 100 µM acetosyringone) to an $OD_{600}$ of 2. After a 2-hour incubation in the dark with gentle shaking, 450 µL of bacterial strains containing the pTRV2-MgChl or pTRV2-FoGS1-MgChl constructs were mixed in equal volume with strains carrying the pTRV1 vector (11). VIGS inoculation of 30-day old *C. roseus* plants, cultivar 'Atlantis Burgundy Halo', was performed by pipetting 10 µL of mixed bacterial strains between the stem and petiole of one leaf grown after the cotyledons. Bacteria were infiltrated by piercing the stem through the bacterial suspension using a ø 0.40 x 25 mm Sterican® needle. The first symptoms of VIGS were observed after 12 days. Silencing of the magnesium chelatase subunit H gene (MgChl) results in yellowing of the leaves (10). Leaf tissue was harvested, and flash frozen in liquid nitrogen 1 week after first observing the yellowing of leaves. *Cr*FoGS1 VIGS was conducted using 6 biological replicates and the experiment was performed twice independently. Frozen VIGS leaf tissue was ground in 2 mL microcentrifuge tubes with tungsten carbide beads (*Qiagen*) using a TissueLyser II homogenizer (*Qiagen*). Metabolites from approximately 10 mg of ground tissue were extracted in 1:30 weight/volume methanol with 2 µM ajmaline as an internal standard. Extracts were vortexed and sonicated for 15 min at room temperature, then leaf tissue was pelleted by centrifugation at 14,000 rpm for 10 min. Supernatant was diluted 300-fold in 100 % methanol and filtered through 0.45 µm low-binding hydrophilic polytetrafluoroethylene spin-filter plates (*Millipore*) and directly analyzed by LC-MS. Transcript abundance was determined by transcriptomics. RNA was isolated from ground VIGS tissue using a RNeasy Mini Kit (*Qiagen*) according to the manufacturer's instructions with an added on-column DNAse digestion. Extracted RNA from all 6 replicates was sent to Novogene for RNA sequencing.

**Recombinant protein expression and purification**

*Cr*SGD, *Cr*GS1, *Cr*FoGS1, *Ti*SGD, *Ti*GS, *Ti*FoGS1, *Ti*FoGS2, *Ti*FoGS3, and *Ti*FoGS1 variants cloned into pOPINF expression vectors with N-terminal $His^6$ tags were recombinantly expressed using *Escherichia coli* BL21 (DE3) cells. Starter cultures were grown overnight in 100 mL LB media at 37 °C with 200 rpm shaking, and subsequently used to inoculate 1 L cultures of LB media. All LB media were supplemented with 100 µg/mL carbenicillin. Cultures were grown at 37 °C with 200 rpm shaking until an $OD_{600}$ value of 0.6-0.8 and cooled for 30 min on ice. Recombinant protein expression was then induced by adding isopropyl β-D-1-thiogalactopyranoside at a final concentration of 1 mM, and cultures were incubated at 18 °C for 12-18 hours with 200 rpm shaking. Cells were harvested by centrifugation at 3200 x *g* at 4 °C for 15 min and cell pellets resuspended in buffer A1 (50 mM Tris-HCl pH 8, 50 mM glycine, 500 mM NaCl, 10 mM imidazole) with lysozyme and EDTA-free protease inhibitor cocktail (*Roche Diagnostics Ltd*.). Resuspended pellets were lysed by sonication and cell debris was subsequently removed by centrifugation at 35,000 x *g* at 4 °C for 30 min. Clarified lysate was incubated with 0.4 mL of Ni/NTA resin (*Qiagen*) for 45 min at 4 °C with gentle shaking. Ni/NTA resin was then washed 3 times with 10 mL of buffer A1 and eluted with 2 consecutive additions of elution buffer (A1 with 250mM imidazole). Imidazole concentration was reduced to less than 1 mM and buffer exchanged into buffer A4 (20 mM HEPES pH 7.5, 150 mM NaCl) by ultrafiltration using 30 kDa molecular weight cut-off centrifugal filters (Merk) Protein purity was assessed by SDS-PAGE (Figure S26). Protein concentration was measured by absorbance at 280 nm based on theoretical extinction coefficients (12). Final concentrated protein was flash frozen in liquid nitrogen at stored at –70 °C.

**Chemoenzymatic synthesis of cathenamine**

Cathenamine was synthesized chemoenzymatically from strictosidine suing *C. roseus* SGD. A total of 5.3 mg of strictosidine was deglycosylated in 25 parallel reactions, each containing 40 nM SGD and 400 µM strictosidine in 50 mM HEPES pH 7.5 in a volume of 2 mL. The reaction was carried out at 30 °C for 4.5 hours and monitored by LC-MS. Pooled reactions were basified with 50 mL of sodium carbonate to adjust to pH 10–11 and extracted 3 times with ethyl acetate (50 mL). The combined organic layer was dried over anhydrous sodium sulphate, filtered, and volatiles were evaporated *in vacuo*. The resulting crude cathenamine was redissolved in 4 mL of 100% methanol and purified using a 1290 Infinity II (*Agillent*) HPLC and 1290 Prep fraction collector. Samples were separated on an Waters XBridge BEH C18 OBD Prep Column (10mm x 250 mm, 5 µm, 130 Å) using mobile phase A (5 mM ammonium acetate in water) and B (pure methanol). The chromatographic method is as follows: 50% B for 3 min, followed by a 50 to 100 % B linear gradient over 20 min, 100 % B for 7 min, 100 to 50 % B in 0.1 min, then a final re-equilibration at 50% B for 4.9 min. Samples were filtered using a 0.22 µm PFTE syringe filter and purified through consecutive injections of 600 µL. Fractions were collected based on 30 sec time slices and analytes detected by UV at 270 nm and 254 nm. The presence of cathenamine (*m/z 351*) was verified on an Expression-L compact mass spectrometer (*Advion Interchim Scientific*) using a mobile phase of acetonitrile/water (70/30) and 0.1 % formic acid. Fractions were pooled, volatiles were evaporated *in vacuo*, then the samples were lyophilized. The purified product (1.23 mg, 23.2% yield) was confirmed to be cathenamine by NMR (Figure S6).

**NMR measurements**

NMR measurements were carried out on a 500 MHz Bruker Avance III HD spectrometer (Bruker Biospin GmbH, Rheinstetten, Germany) using standard pulse sequences as implemented in Bruker Topspin ver. 3.6.1. 500 MHz NMR equips with a TCI cryoprobe. Chemical shifts were referenced to the residual solvent signals of $CDCl_3$ (□$_H$ 7.26/□$_C$ 77.16) (13). All spectra were recorded at 298 K. NMR spectra were processed by Bruker Topspin ver. 3.6.1. (13).

**Sodium borohydride reduction of strictosidine aglycone**

Strictosidine aglycone was prepared in 50 µL reactions by degylcosylating 200 µM of strictosidine by 20nM SGD in 50 mM HEPES pH 7.5 at 30 °C for 2 hours. Sodium borohydride was added to a final concentration of 10 mM and reactions were incubated at room temperature for 30 min. Reactions were basified with 200 µL of sodium carbonate to adjust to pH 10–11 and extracted twice with ethyl acetate (500 µL). Volatile solvents were evaporated on a flow of nitrogen gas and re-suspended in 200 µL 100% methanol. Samples were filtered using a 0.22 µm PFTE syringe filter and analyzed using the same LC-MS method as *N. benthamiana* and *in vitro* assays.

**Subcellular localization in *N. benthamiana* leaves**

The subcellular localization of SGD, FoGS1, GS, and HYS from *C. roseus* and *T. iboga* was determined in *N. benthamiana* leaves. eGFP tagged (N-terminal) genes of interest (pCambia) were overexpressed in *N. benthamiana* with untagged mCherry (3omega1) as a nucleocytosolic marker and a mCherry-tagged nuclear marker (14). Pair-wise co-expressions of eGFP tagged (N-terminal) and mRFP1 tagged (C-terminal) genes of interest were also conducted. Leaf discs were excised from *N. benthamiana* leaves 3-days post agrobacterium infiltration and mounted in water. Imaging was conducted on a cLSM

880 (Zeiss, Oberkochen, Germany) equipped with a C-Apochromat 40x/1.2 water objective. GFP was excited using an argon laser at a wavelength of 488 nm with 1-10 % transmission and 700 PMT gain. RFP was excited using a helium-neon laser at a wavelength of 543 nm with 10-20 % transmission and 700 PMT gain. Emission was detected between 490 and 550 nm for GFP and between 550 and 650 nm for RFP. Transmitted light was measured in the GFP channel with a T-PMT gain of 600. Images were acquired using a pixel dwell time of approximately 1 µ-second with 8-fold line averaging, and a Pinhole size of 1 Airy Unit. Image contrast and brightness were edited in Image J (15).

**Synteny analysis**

Genome sequences (FASTA), gene annotation files (GFF), and gene coding sequences (CDS) were processed using MCscan (Python version) from the JCVI utility libraries (16); https://github.com/tanghaibao/jcvi). Pairwise homolog searches were performed using **LAST** ((17); https://gitlab.com/mcfrith/last). The resulting LAST hits were filtered to remove low-quality alignments and collapsed to eliminate tandem duplicates before MCscan anchor identification. The filtered anchors were subsequently clustered into collinear blocks using default MCscan settings. Microsynteny plots were generated with the synteny function under default visualization parameters.

**Liquid Chromatography – Mass Spectrometry (LC-MS) methods**

Data acquisition from *N. benthamiana* and *in vitro* assays were performed on an UltiMate 3000 ultrahigh performance system (*Thermo Fischer*) coupled to an Impact II ultra-high-resolution quadrupole-time-of-flight mass spectrometer (*Bruker*). Analytes were separated on an XBridge BEH C18 colunm (4.6 x 100 mm, 2.5 µm) operated at 50 °C. Samples were injected (2 µL) and run at a flow rate of 0.65 mL/min, using mobile phase A (5 mM ammonium acetate in water) and B (pure methanol). The chromatographic method is as follows: 50 % B for 1 min, followed by a 50 to 95 % B linear gradient over 7.5 min, 95 to 50 % B in 0.1 min, 95 % B for 0.5 min, and a final re-equilibration at 50 % B for 1.9 min. Mass spectrometry data acquisition was performed in positive electrospray ionization mode with an end-plate offset of 500 V, capillary voltage of 3500 V, nebulizer pressure of 2.5 Bar, drying nitrogen flow at 11 L/min, and a dry gas temperature of 250 °C. MS data was recorded in the *m/z* range of 80 to 1000 at 12 Hz. Tandem spectrometry data was acquired in data-dependent mode with a threshold of 400 counts to trigger fragmentation using stepping collision energy of 20 to 50 eV. Total cycle time was 0.5 sec. Each run was externally calibrated with a solution of sodium formate-isopropanol loaded at 0.18 mL/min. The first 3 min and last 2 min were directed to waste. Data was analyzed using Bruker Compass Hystar v.6.2 software.

Data acquisition for VIGS was performed on an a Vanquish ultrahigh performance system (*Thermo Fisher*) coupled to a Q-Exactive Plus orbitrap mass spectrometer (*Thermo Fisher*) using Xcalibur software version 4.7.69.37 (*Thermo Fisher*). Samples were separated on a Waters Acquity UPLC® BEH C18 (2.1 x 50 mm, 1.7 µm) column operated at 40 °C, based on a previously described method (3). Samples were injected (2 µL) and run at a flow rate of 0.6 mL/min, using mobile phase A (0.1 % formic acid in water) and B (acetonitrile). The chromatographic method is as follows: 10 % B for 0.4 min, followed by a 20 to 42 % B linear gradient over 2.8 min, 42 to 100 % B over 0.05 min, 100 % B for 0.5 min, 100 to 10 %B in 0.05 min, and a final re-equilibration at 10 % B for 1.2 min. Mass spectrometry data acquisition was performed in positive mode using heated electrospray ionization with sheath gas flow rate of 55, auxiliary gas flow rate of 15, sweep gas flow rate of 3, spray voltage of 3.5 kV, capillary temperature of 275 °C, auxiliary gas heater temperature of 450 °C, and S-lens RF level of 50. MS data was acquired in full-scan MS mode in the *m/z* range of 120 to 1000 with a resolution of 70000-FWHM, chromatogram peak width of 7 sec, automatic gain control target of $5 \times 10^5$, using a centroid spectrum format. The first 0.3 min and last 1.2 min were directed to waste. The instrument was calibrated using the Pierce LTQ Velos ESI positive ion calibration solution. Data was processed in Xcalibur software version 4.7.69.37 (*Thermo Fisher*) using the retention times of authentic standards, and integrated peak area were normalized to the internal standard (ajmaline).

Alternate LC-MS method for detection of cathenamine, modified from a previous report (18). Briefly, cathenamine was detected on an UltiMate 3000 ultrahigh performance system (*Thermo Fischer*) coupled to an Impact II ultra-high-resolution quadrupole-time-of-flight mass spectrometer (*Bruker*). Separation was achieved using a Phenomenex Kinetex XB C18 column (100 x 2.1 mm, 2.6 µm; 100 Å) operated at 40 °C with mobile phase A (0.1 % formic acid in water) and mobile phase B (acetonitrile), using a flow rate of 0.6 mL/min and 2 µL sample injections. The chromatographic method is as follows: 10% B for 1 min, followed by a 10 to 30 % B linear gradient over 5 min, 90 % B for 1.5 min, then a final re-equilibration at 10% B for 2.5 min. Mass spectrometry data acquisition was performed in positive electrospray ionization mode with an end-plate offset of 500 V, capillary voltage of 3500 V, nebulizer pressure of 2.5 Bar, drying nitrogen flow at 11 L/min, and a dry gas temperature of 250 °C. MS data was recorded in the *m/z* range of 80 to 1000 at 12 Hz. Tandem spectrometry data was acquired in data-dependent mode with a threshold of 400 counts to trigger fragmentation using stepping collision energy of 20 to 50 eV. Total cycle time was 0.5 sec. Each run was externally calibrated with a solution of sodium formate-isopropanol loaded at 0.18 mL/min. The first 3 min and last 2 min were directed to waste. Data was analyzed using Bruker Compass Hystar v.6.2 software.

**Protein structural modelling**

Structural predictions for the TiFoGS-GS heterodimer with NADPH cofactor and coordinated zinc ions were generated using AlphaFold3 (19). Molecular docking was performed using autodock4$_{Zn}$ (20) using an initial prediction of 19*E*-dehydrogeissoschizine three-dimensional structure from Chem3D. The docking pose with the lowest predicted binding energy was selected.

**Supplemental Figures**

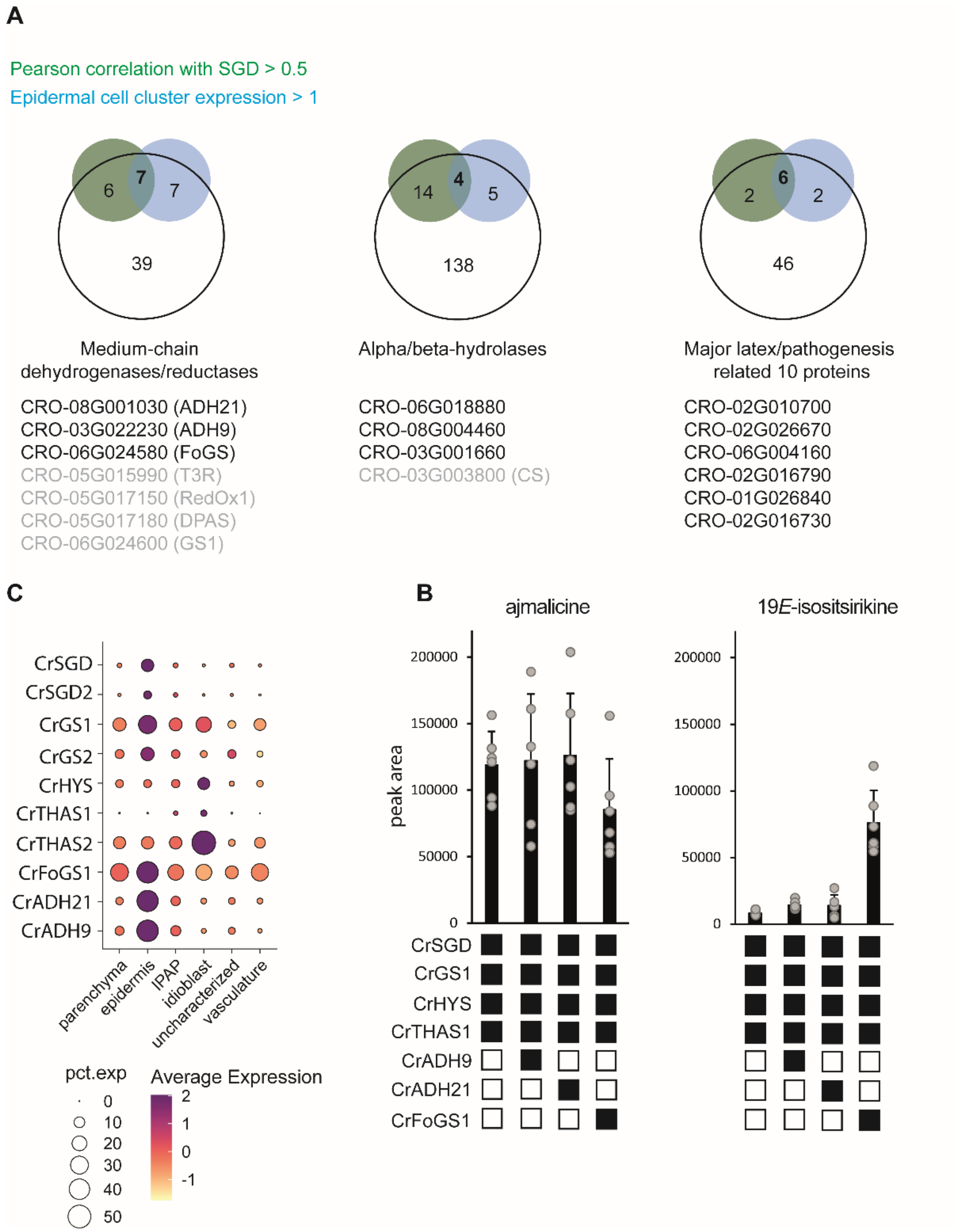


**Figure S1: Candidate selection and screening in *N. benthamiana*.** **A** *C. roseus* single cell transcriptomics data for candidates annotated as medium chain dehydrogenases along with select MIA biosynthetic genes. Single cell transcriptomic expression levels (3) are represented by a dotplot where circle size corresponds to percent expression and circle colour corresponds to average expression. Although HYS is predominantly expressed in idioblast cells, a different dataset (21) shows HYS-expression in both epidermal and idioblast cells. **B** Screening of candidate medium-chain dehydrogenases/reductases in *N. benthamiana*. Each treatment is the average of 6 biological replicates. Values for individual replicates are shown as grey circles and averages are shown with error bars. SGD, strictosidine β-glucosidase; GS, geissoschizine synthase; FoGS, facilitator of geissoschizine synthase;

HYS, heteroyohimbine synthase; THAS, tetrahydroalstonine synthase; ADH, alcohol dehydrogenase. **C** Ven diagram visualization of candidate selection. Candidate medium-chain dehydrogenase/reductases, major latex/pathogenesis related 10 proteins, and alpha/beta-hydrolases were filtered based on Pearson correlation with SGD cell type expression (green) and level of expression in epidermal cell clusters (blue). Only candidates that met the cutoff values of 0.5 (Pearson correlation) and FPKM values above 1 (epidermal cell cluster expression) were selected for screening in *N. benthamiana*. Candidate gene IDs are listed below Ven diagrams. Grey IDs correspond to known biosynthetic genes and were not screened. ADH; alcohol dehydrogenase; SGD, strictosidine β-glucosidase; GS, geissoschizine synthase; FoGS1, facilitator of geissoschizine synthase; HYS, heteroyohimbine synthase; THAS, tetrahydroalstonine synthase; RedOx1, reductive oxidative enzyme 1; DPAS, dehydroprecondylocarpine synthase; T3R; tabersonine 3-reductase; CS, catharanthine synthase.

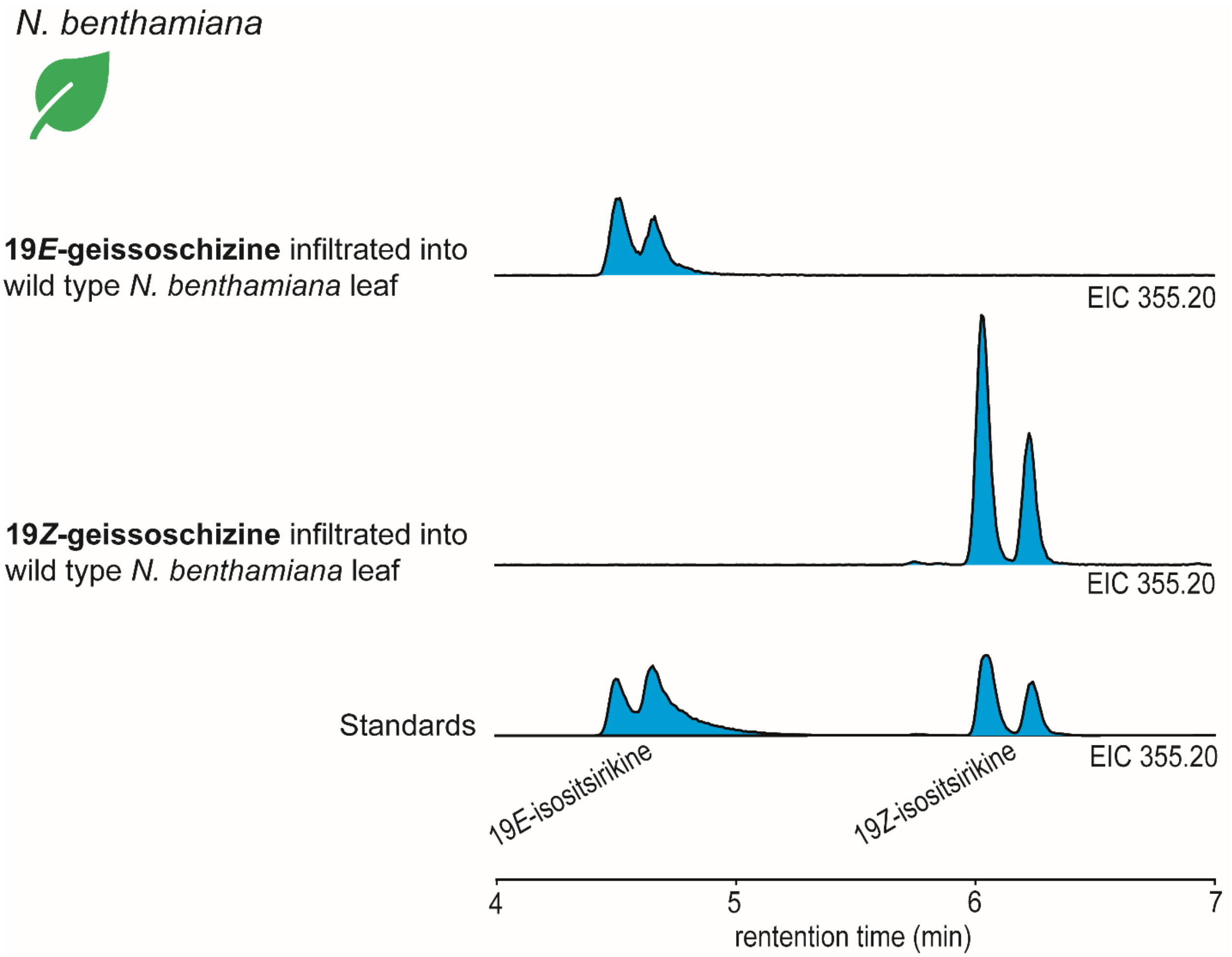


**Figure S2: Reduction of 19*E*- and 19*Z*-geissoschizine by endogenous *N. benthamiana* enzyme(s).** Representative extracted ion chromatograms (EIC) for *m/z 355.20* of wild type *N. benthamiana* leaves infiltrated with 19*E*- or 19*Z*-geissoschizine. Standards for 19*E*- and 19*Z*-isositsirikine are shown.

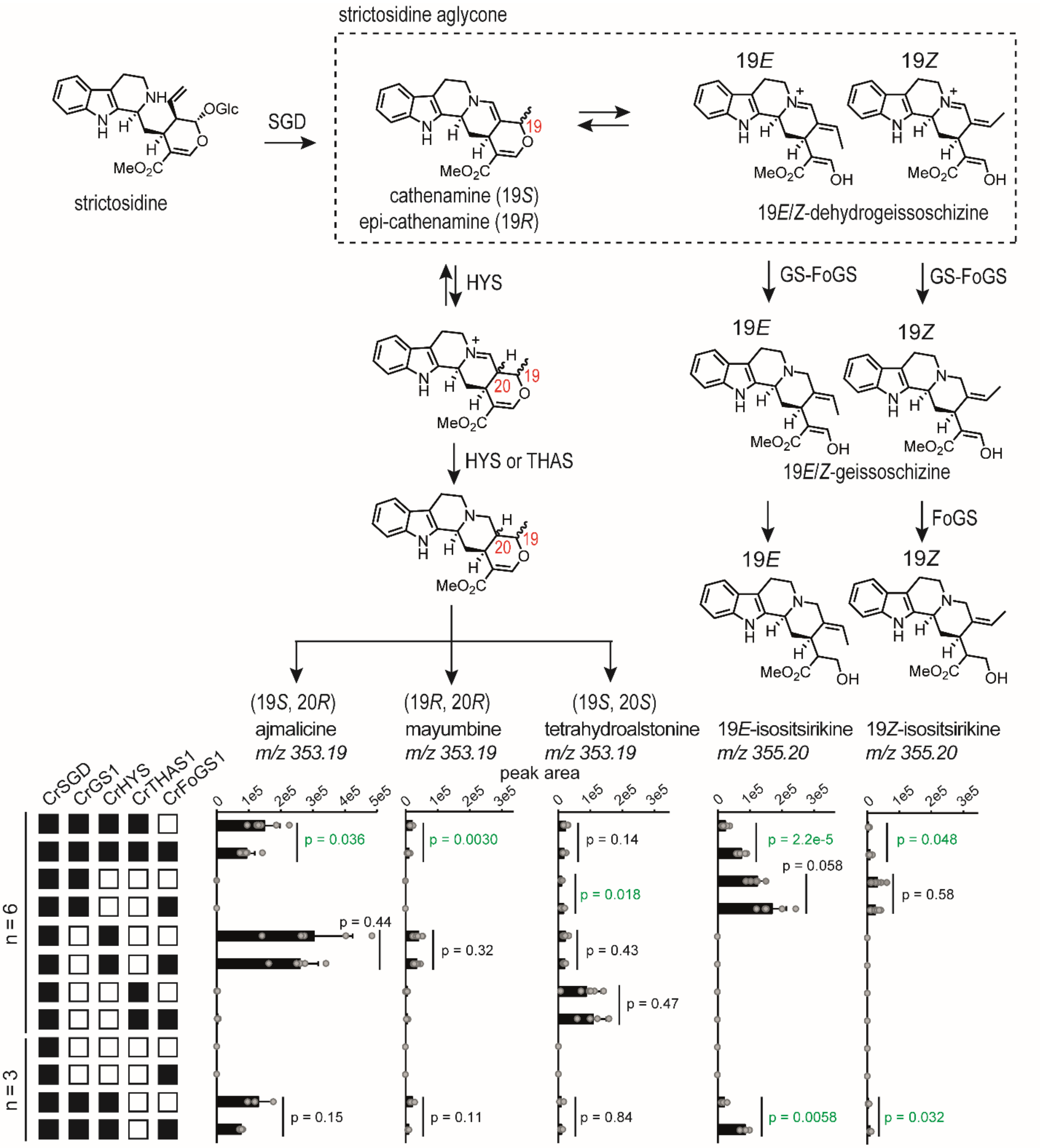


**Figure S3: Deconvolution of competition assays with *Cr*GS1, *Cr*HYS, and *Cr*THAS1 and with and without *Cr*FoGS1 in *N. benthamiana*. A** Overview of the chemical conversions, and corresponding enzymes, of strictosidine to the detected products in *N. benthamiana* assays. **B** Deconvolution of competitive GS1, HYS, THAS1 assays with and without FoGS1, in *N. benthamiana*. Each treatment is the average of 3 or 6 biological replicates. Values for individual replicates are shown as grey circles and averages are shown with error bars and *p*-values from student's t-tests. *p*-values below 0.05 as shown in green. SGD, strictosidine β-glucosidase; GS, geissoschizine synthase; FoGS, facilitator of geissoschizine synthase; HYS, heteroyohimbine synthase; THAS, tetrahydroalstonine synthase.

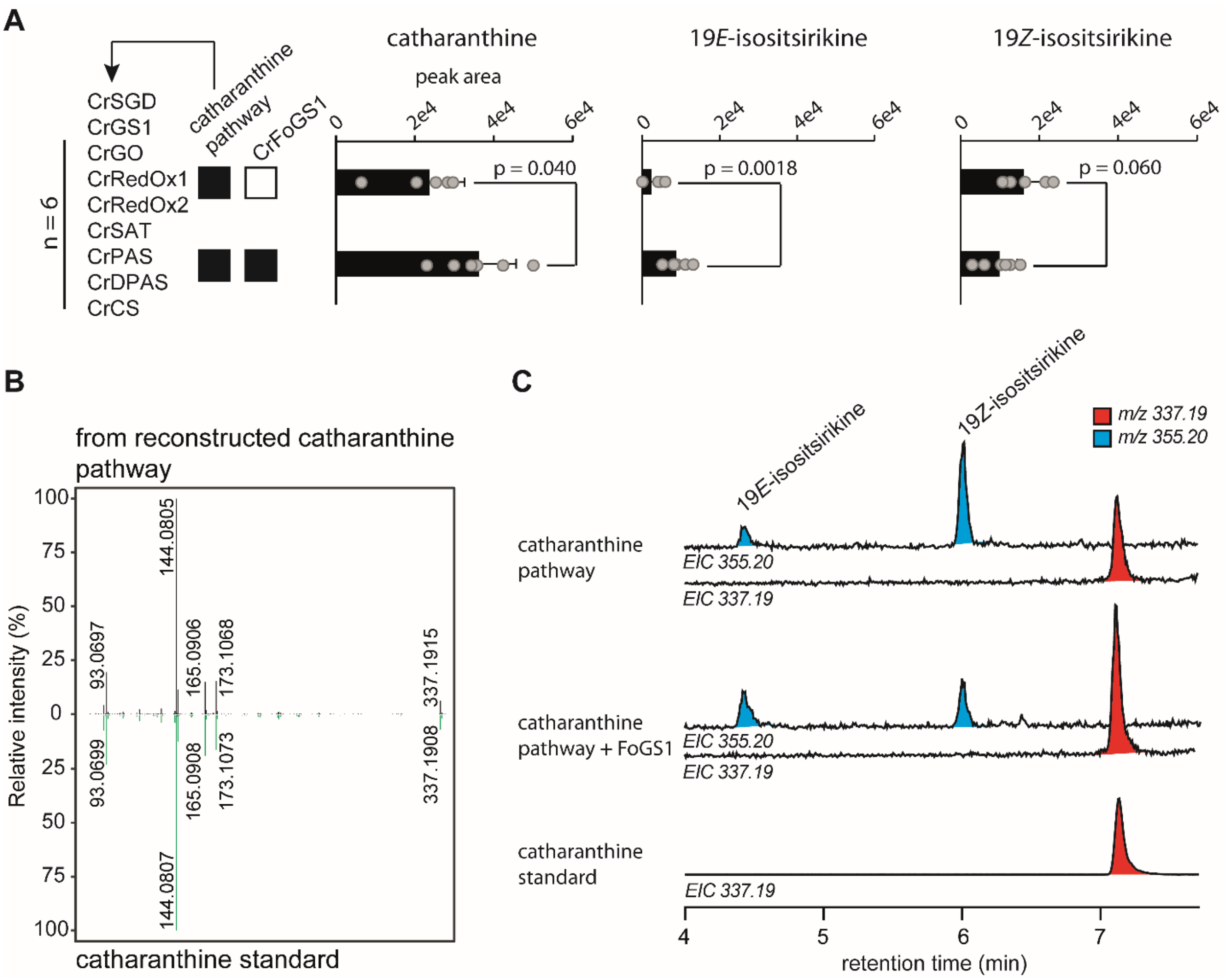


**Figure S4: Reconstitution of the biosynthetic pathway from strictosidine to catharanthine (*C. roseus*) biosynthetic pathway in *N. benthamiana*. A** Peak areas from 6 replicates presented in Figure 2D. Values for individual replicates are shown as grey circles and averages are shown with error bars and *p*-values from student's t-tests. SGD, strictosidine β-glucosidase; GS, geissoschizine synthase; FoGS1, facilitator of geissoschizine synthase; GO, geissoschizine oxidase; RedOx1, reductive oxidative enzyme 1; RedOx2, reductive oxidative enzyme 2; SAT, stemmadenine acetyl transferase; PAS, precondylocarpine acetate synthase; DPAS, dehydroprecondylocarpine synthase; CS, catharanthine synthase. **B** LC-MS/MS spectra of a catharanthine standard (green) and catharanthine from pathway reconstitution in *N. benthamiana* (black). **C** Representative extracted ion chromatograms (EIC) of *m/z 337.19* (red) and *355.20* (cyan).

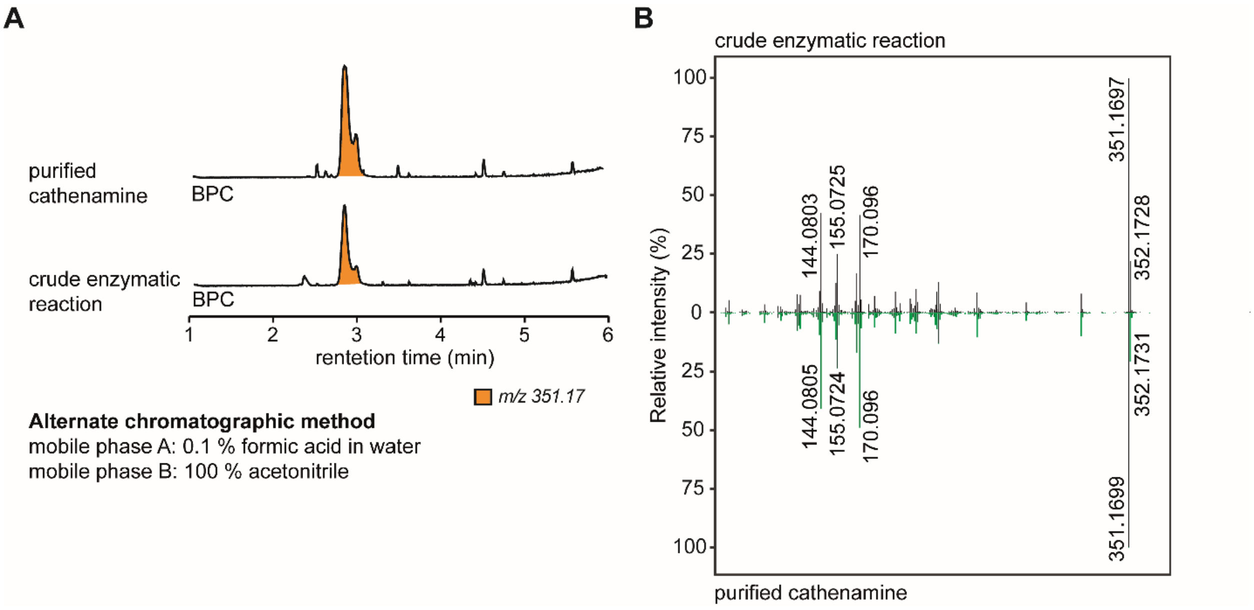


**Figure S5: Purification of cathenamine. A** Base peak chromatograms (BPC) of the crude product of enzymatic *Cr*SGD deglycosylation of strictosidine and the final purified cathenamine after reaction upscaling and purification. LC-MS analysis was performed using an alternate method to achieve better chromatographic separation, modified from a previous report (18). **B** LC-MS/MS spectra of crude product of enzymatic *Cr*SGD deglycosylation of strictosidine (black) and the final purified cathenamine (green).

**A**

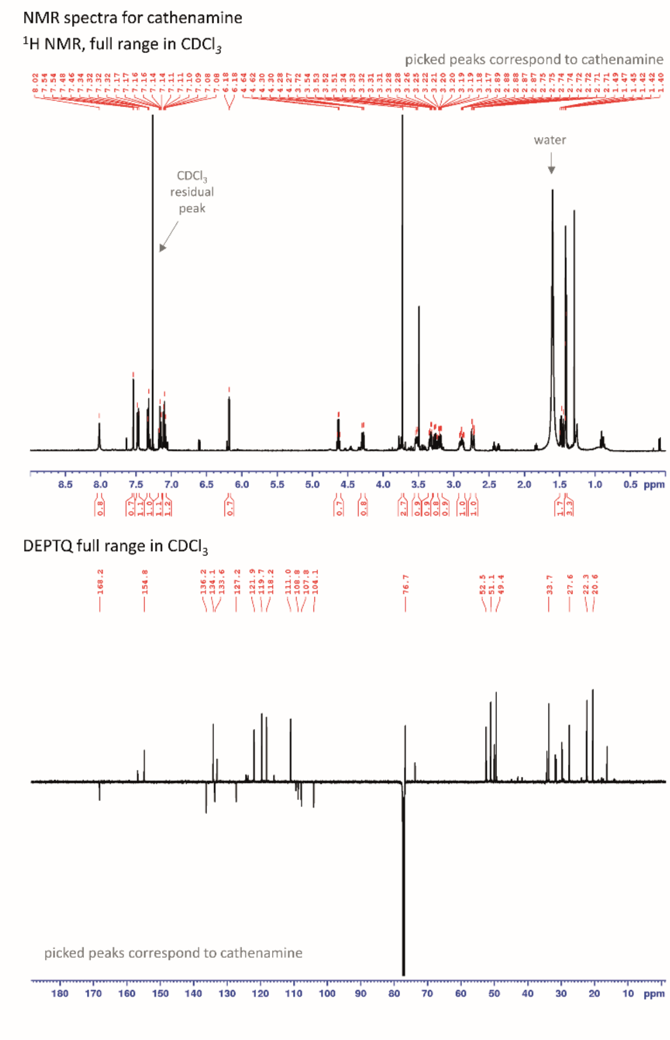


**B**

| pos. | $\delta_H$ | *mult.* | $J_{HH}$ | $\delta_C$ |
|---|---|---|---|---|
| 1(NH) | 8.02 | *brs* | - | - |
| 2 | - | - | - | 133.6 |
| 3α | 4.29 | *dddd* | 11.7/2.4/2.1/2.0 | 52.5 |
| 5α | 3.26 | *ddd* | 12.1/10.4/4.0 | 49.4 |
| 5β | 3.33 | *ddd* | 12.1/5.7/2.3 | 49.4 |
| 6α | 2.73 | *dddd* | 15.0/4.0/2.3/2.0 | 22.3 |
| 6β | 2.89 | *dddd* | 15.0/10.4/5.7/2.1 | 22.3 |
| 7 | - | - | - | 108.8 |
| 8 | - | - | - | 127.2 |
| 9 | 7.47 | *brd* | 7.8 | 118.2 |
| 10 | 7.10 | *ddd* | 7.8/7.2/1.0 | 119.7 |
| 11 | 7.16 | *ddd* | 8.0/7.2/1.1 | 121.9 |
| 12 | 7.33 | *ddd* | 8.0/1.0/0.8 | 111.0 |
| 13 | - | - | - | 136.2 |
| 14α | 3.19 | *ddd* | 12.8/5.8/2.4 | 33.7 |
| 14β | 1.46 | *ddd* | 12.8/11.7/11.3 | 33.7 |
| 15α | 3.52 | *m* | - | 27.6 |
| 16 | - | - | - | 107.8 |
| 17 | 7.54 | *d* | 1.4 | 154.8 |
| 18 | 1.41 | *d* | 6.7 | 20.6 |
| 19β | 4.63 | *q* | 6.7 | 76.7 |
| 20 | - | - | - | 104.1 |
| 21 | 6.18 | *d* | 1.9 | 134.1 |
| 22 | - | - | - | 168.2 |
| OMe | 3.72 | *s* | - | 51.1 |

500 MHz in $CDCl_3$
The chemical shifts agreed well with the published data*.

**C**

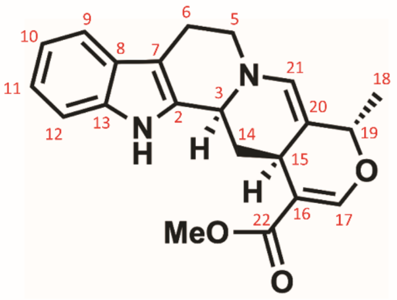


**Figure S6: Cathenamine NMR. A** NMR spectra of cathenamine ($CDCl_3$; 500 MHz for $^1H$, 125 MHz for $^{13}C$, respectively). **B** Peak assignment of cathenamine (13). **C** Numbering of cathenamine.

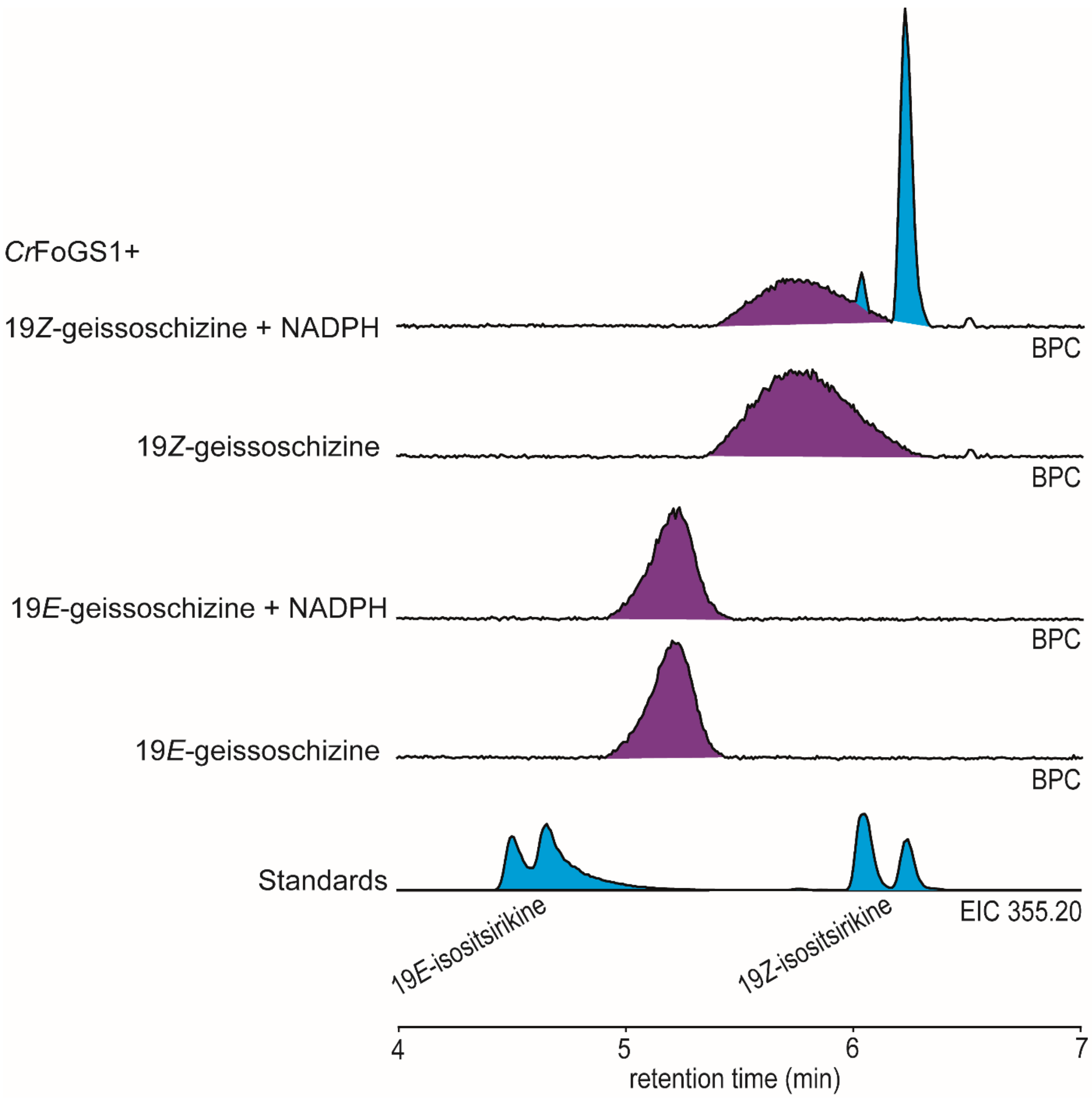


**Figure S7: *In vitro* assays of *Cr*FoGS1 with 19*E*- and 19*Z*-geissoschizine.** Base peak chromatograms (BPC) with *m/z 355.20* peaks highlighted in cyan and m/z *353.19* peaks in purple. *Cr*FoGS1 was assayed with 19*E*- and 19*Z*-geissoschizine with and without NADPH.

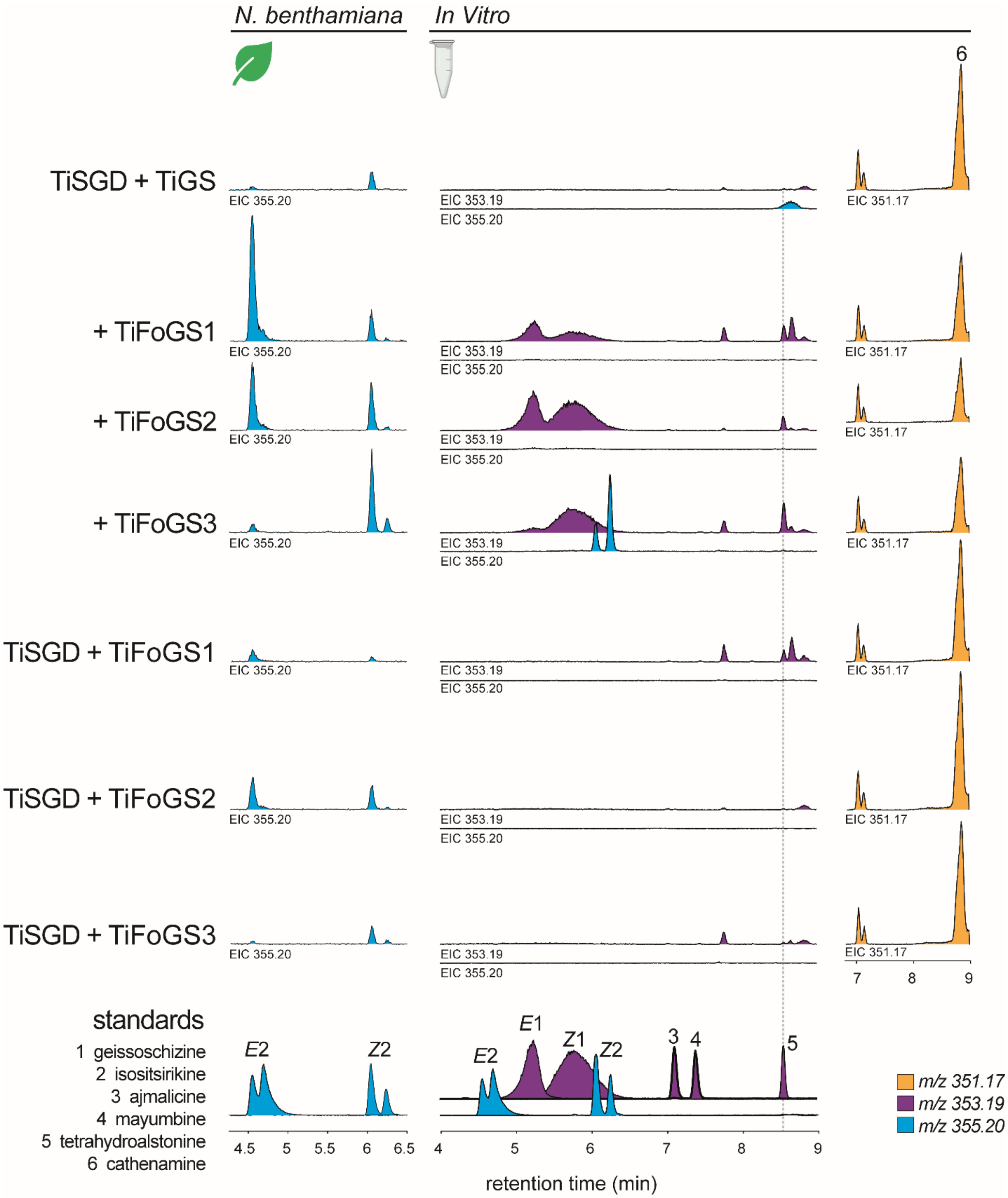


**Figure S8: *N. benthamiana* and *in vitro* characterization of *Ti*FoGS1, *Ti*FoGS2, and *Ti*FoGS3.** TiFoGS orthologues assayed (*in vitro*) or co-expressed (*N. benthamiana*) with TiGS and *Ti*SGD using strictosidine as a substrate. Extracted ion chromatograms (EIC) are shown for *m/z* values of *353.19* (purple), *355.20* (cyan), and *351.17* (orange). Standards for 19*E*/*Z*-geissoschizine (1), 19*E*/*Z*-isositsirikine (2), ajmalicine (3), mayumbine (4), and tetrahydroalstonine (5) are shown below.

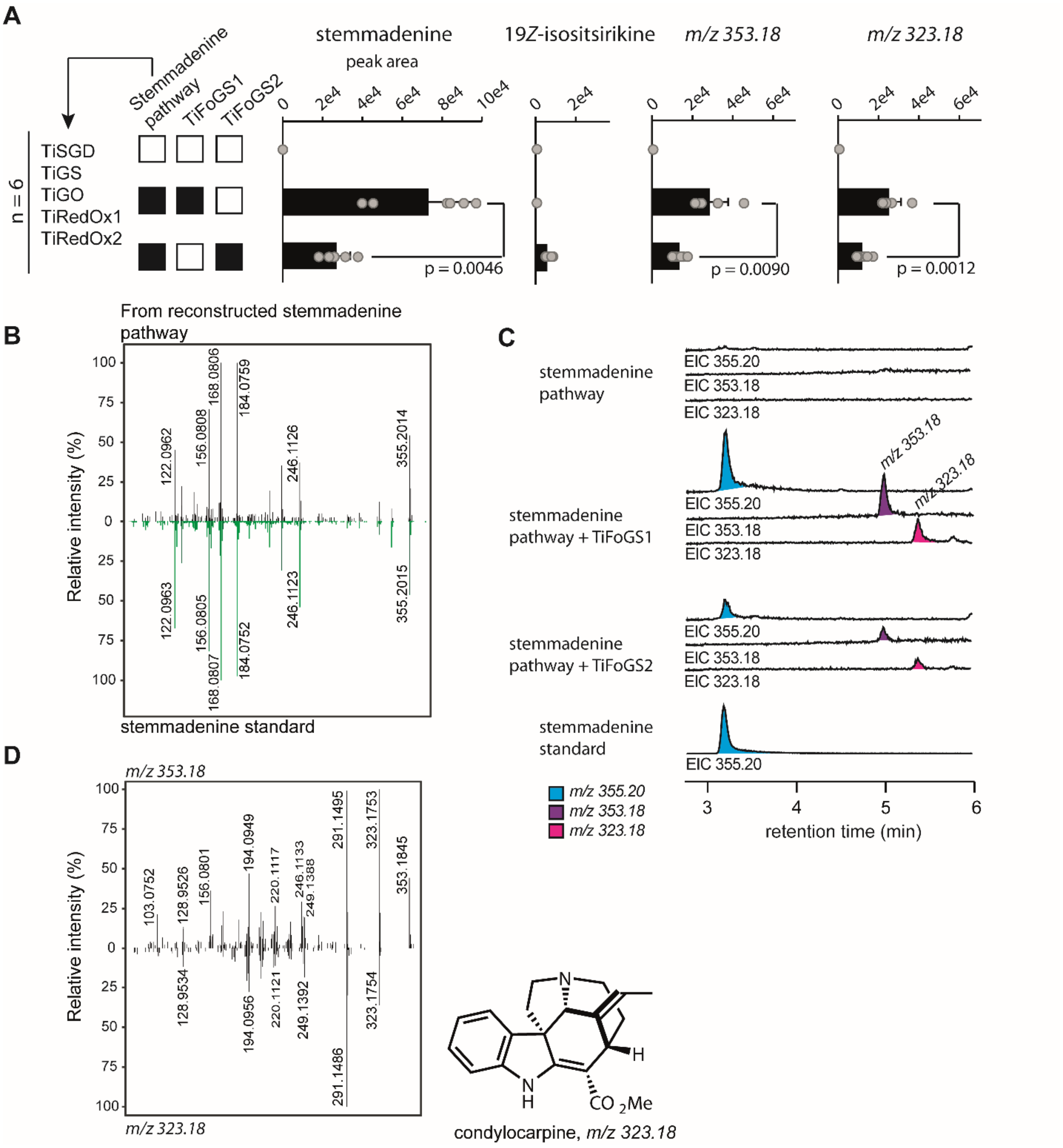


**Figure S9: Reconstitution of the *T. iboga* strictosidine to stemmadenine biosynthetic pathway in *N. benthamiana*.** **A** Peak areas from 6 replicates presented in Figure 3B. Values for individual replicates are shown as grey circles and averages are shown error bars and *p*-values from student's t-tests. SGD, strictosidine β-glucosidase; GS, geissoschizine synthase; FoGS1, facilitator of geissoschizine synthase; GO, geissoschizine oxidase; RedOx1, reductive oxidative enzyme 1; RedOx2, reductive oxidative enzyme 2. **B** MS/MS spectra of a stemmadenine standard (green) and stemmadenine from pathway reconstitution in *N. benthamiana* (black). **C** Representative extracted ion chromatograms (EIC) of *m/z 355.20* (cyan), *353.18* (purple), and *323.18* (pink). **D** The MS/MS spectrum of the *m/z 323.18* shunt product is near identical to that reported for condylocarpine (2), which was shown to be a product of endogenous *N. benthamiana* conversion of stemmadenine. The *m/z 353.18* shunt product has not been previously reported, but is likely condylocarpine based on the MS/MS spectrum.

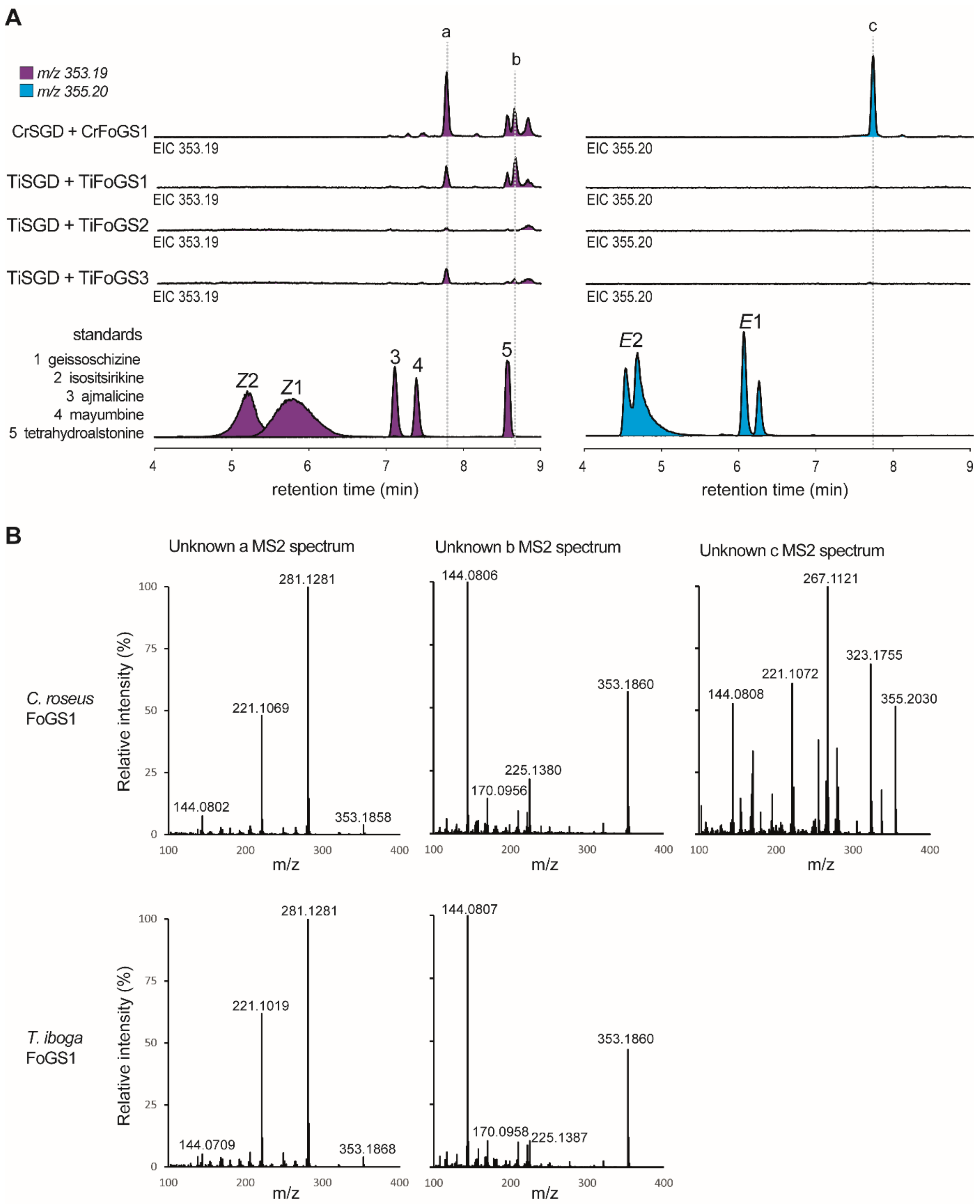


**Figure S10: Uncharacterized minor products produced by *Cr*FoGS and *Ti*FoGS. A** *In vitro* assays of SGD and FoGS orthologues with strictosidine. Extracted ion chromatograms (EIC) for *m/z 353.19* (purple) and *355.20* (cyan) are shown. Major unknown peaks are marked. **B** LC-MS/MS spectra for unknown a, b, and c.

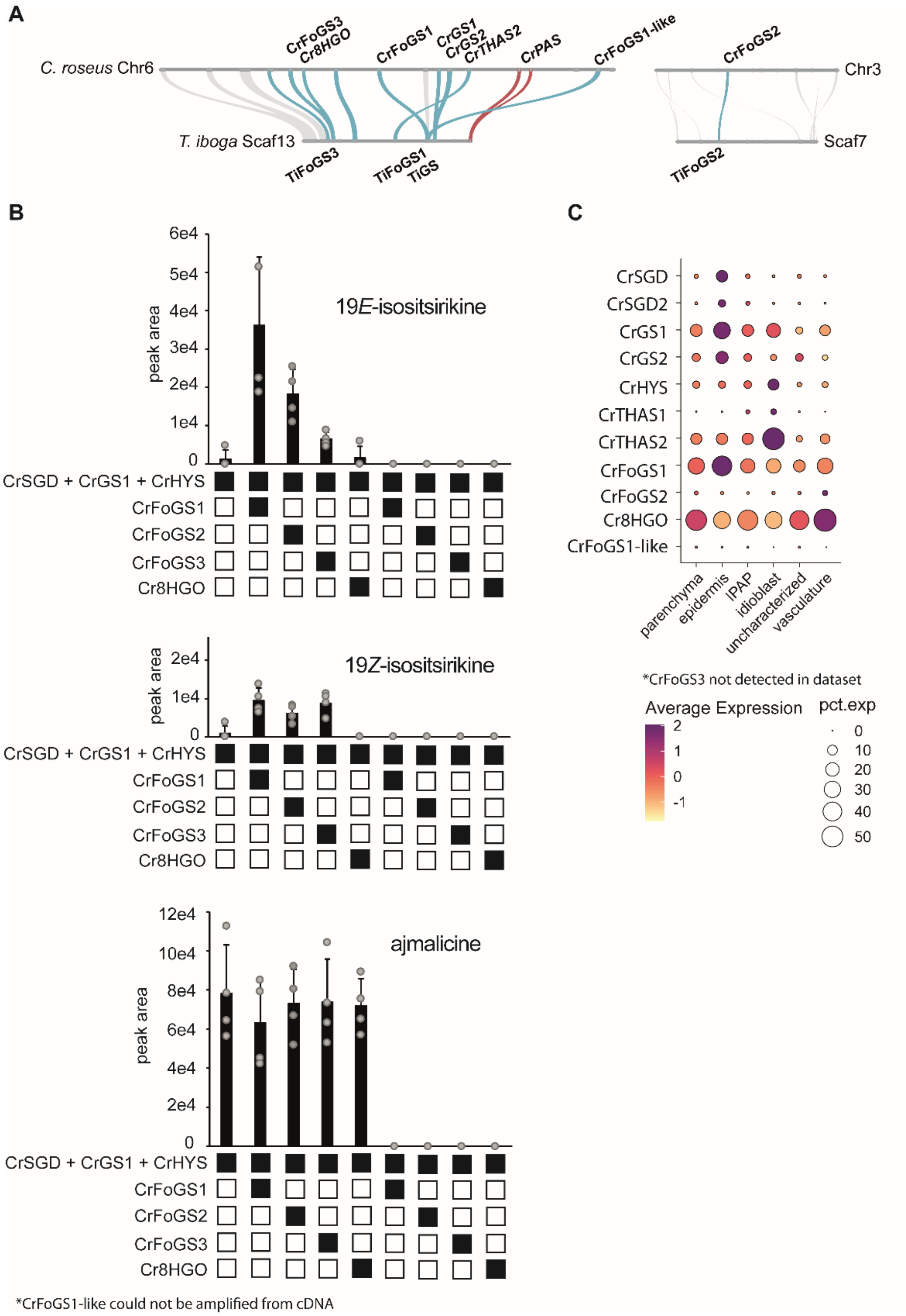


**Figure S11: Characterization of FoGS syntenologues from *C. roseus* in *N. benthamiana*. A** Synteny analysis of *C. roseus* and *T. iboga* FoGS orthologues. **B** *N. benthamiana* assays of *C. roseus* syntenologues of *Ti*FoGS2 and *Ti*FoGS3. Candidates were co-expressed with *Cr*SGD, *Cr*GS1, and *Cr*HYS and assayed with strictosidine. Values for individual replicates are shown as grey circles and averages as black bars with error bars. Four biological replicates were used. **C** Single cell transcriptomics for selected *C. roseus* genes. Single cell transcriptomics are represented by a dotplot where circle size corresponds to percent expression and circle colour corresponds to average expression.

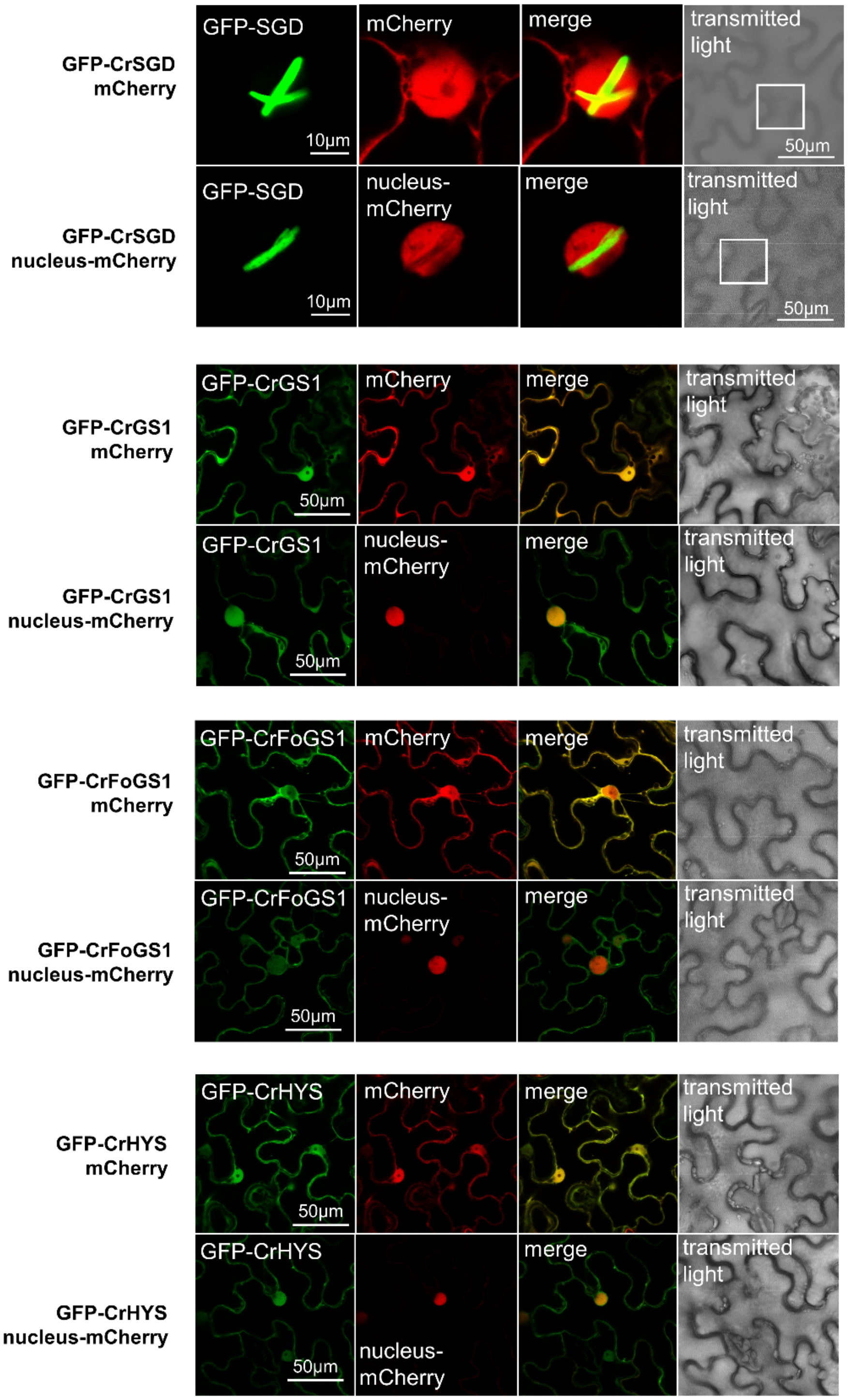


**Figure S12: Subcellular localization of *C. roseus* SGD, GS1, FoGS1, and HYS in *N. benthamiana* leaves.** The proteins of interest were expressed with an N-terminal GFP tag. Co-expression with an mCherry-tagged nuclear marker and nucleocytosolic free-mCherry is shown. GFP fluorescence is shown in green, mCherry in red, and merged GFP-mCherry in yellow.

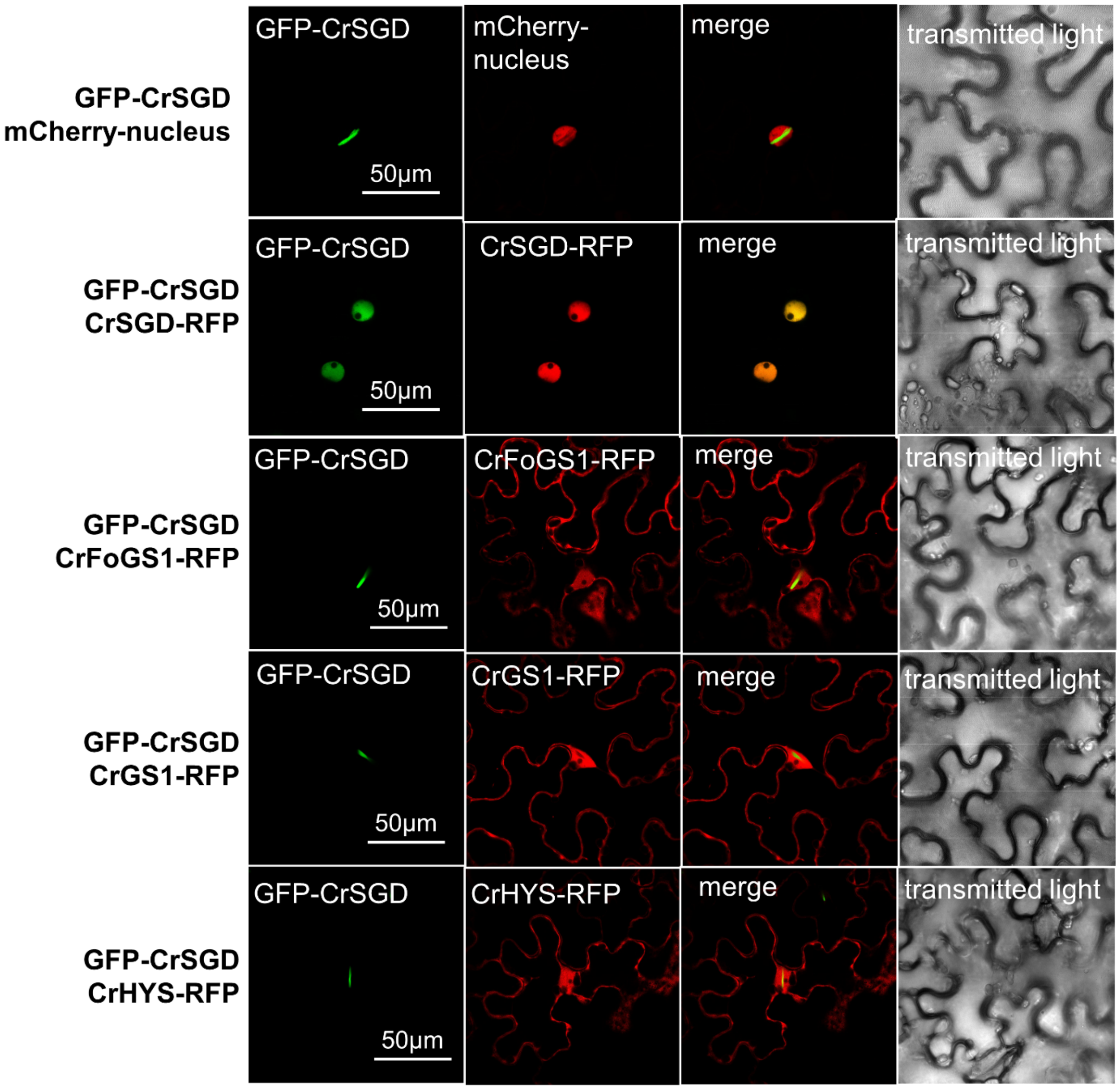


**Figure S13: Subcellular localization of *C. roseus* SGD when co-expressed with SGD, GS1, FoGS1, or HYS in *N. benthamiana* leaves.** N-terminal GFP tagged *Cr*SGD was co-expressed with C-terminal RFP tagged genes of interest. *Cr*SGD co-expression with an mCherry-tagged nuclear marker is shown for reference. GFP fluorescence is shown in green, RFP/mCherry in red, and merged GFP-RFP/mCherry in yellow.

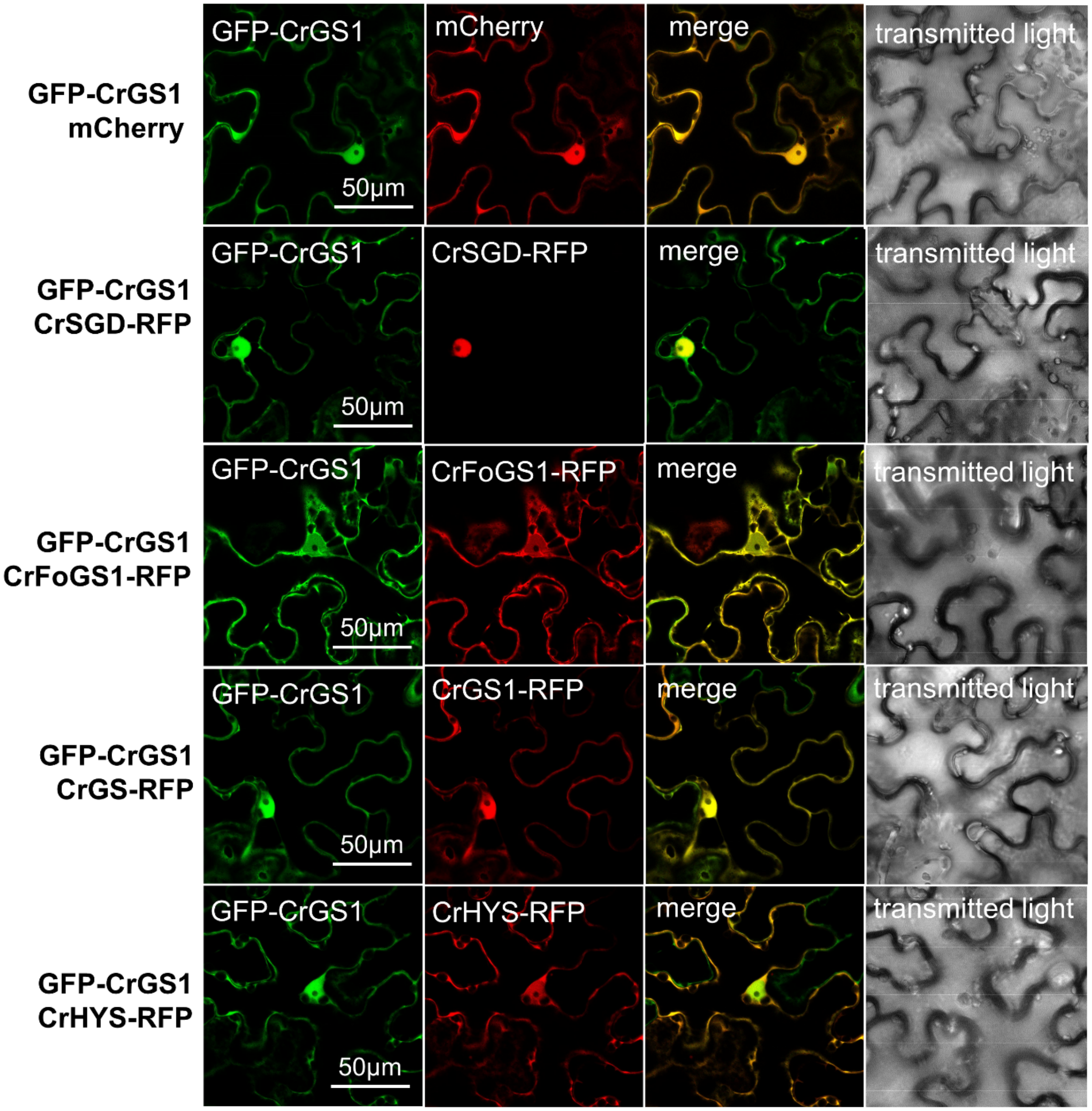


**Figure S14: Subcellular localization of *C. roseus* GS1 when co-expressed with SGD, GS1, FoGS1, or HYS in *N. benthamiana* leaves.** N-terminal GFP tagged *Cr*GS was co-expressed with C-terminal RFP tagged genes of interest. *Cr*GS1 co-expression with nucleocytosolic free-mCherry is shown for reference. GFP fluorescence is shown in green, RFP/mCherry in red, and merged GFP-RFP/mCherry in yellow.

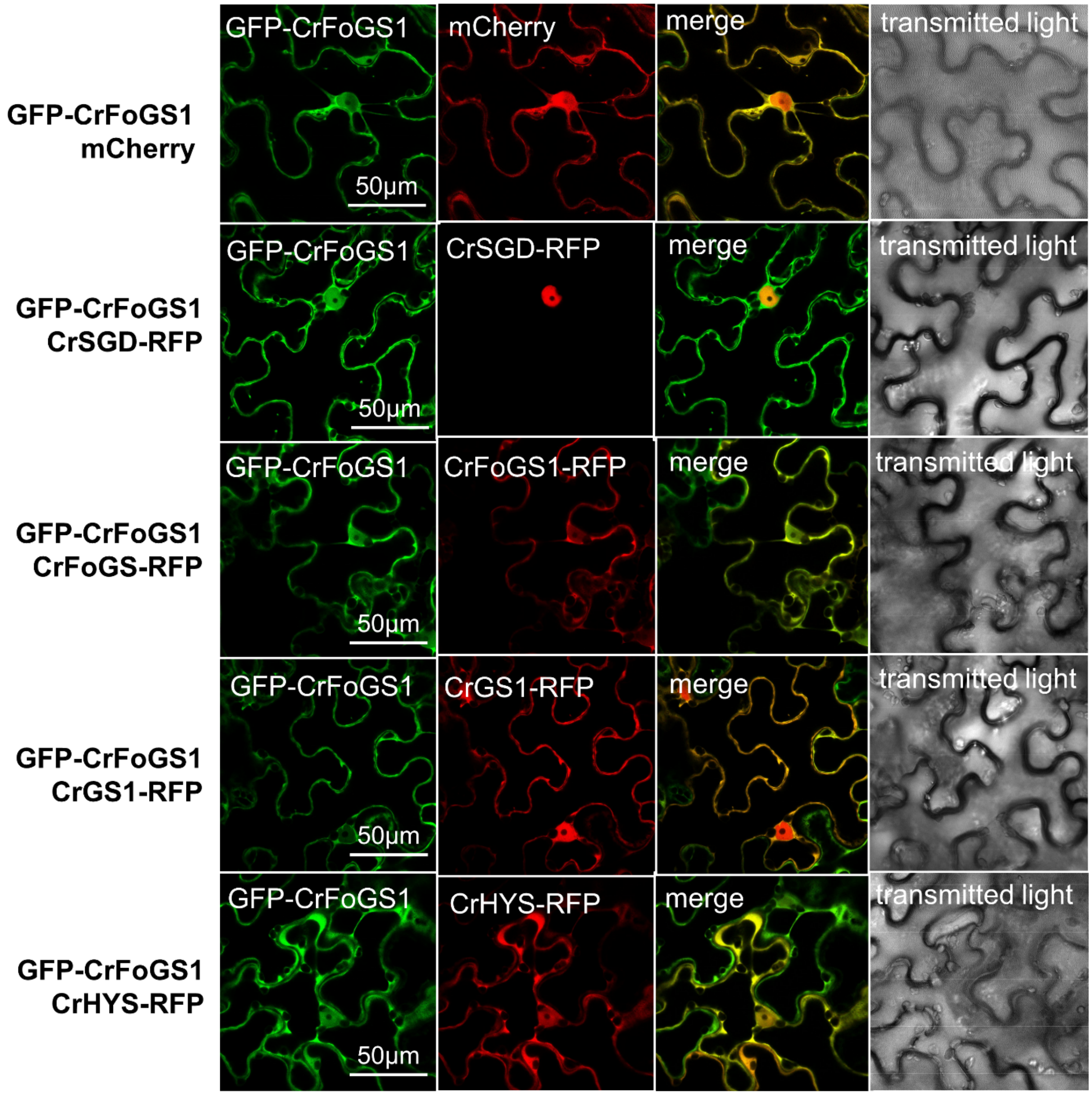


**Figure S15: Subcellular localization of *C. roseus* FoGS1 when co-expressed with SGD, GS1, FoGS1, or HYS in *N. benthamiana* leaves.** N-terminal GFP tagged *Cr*FoGS1 was co-expressed with C-terminal RFP tagged genes of interest. *Cr*FoGS1 co-expression with nucleocytosolic free-mCherry is shown for reference. GFP fluorescence is shown in green, RFP/mCherry in red, and merged GFP-RFP/mCherry in yellow.

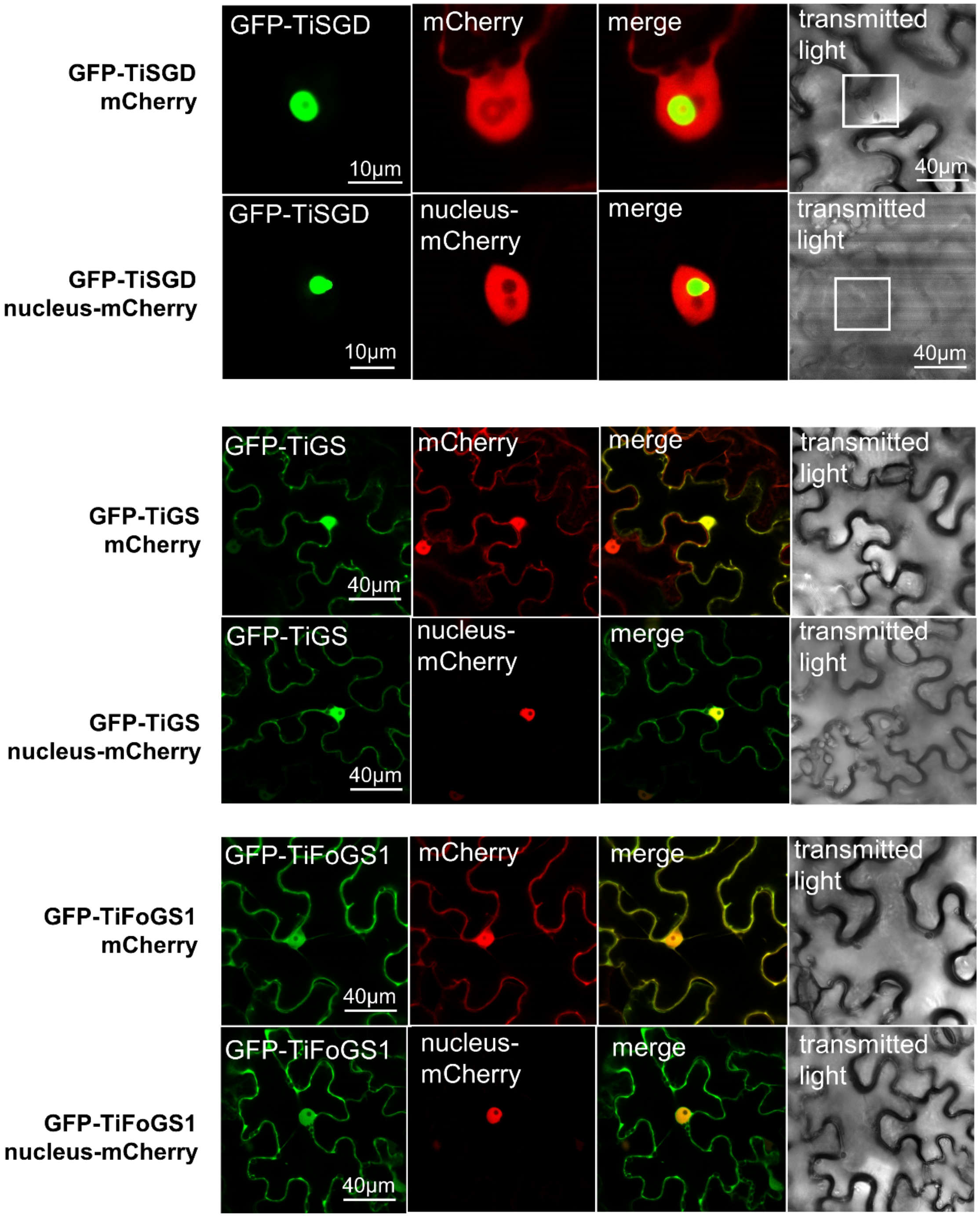


**Figure S16: Subcellular localization of *T. iboga* SGD, GS, and FoGS1 in *N. benthamiana* leaves.** The proteins of interest were expressed with an N-terminal GFP tag. Co-expression with an mCherry-tagged nuclear marker and nucleocytosolic free-mCherry is shown. GFP fluorescence is shown in green, mCherry in red, and merged GFP-mCherry in yellow. Results presented here are consistent with previous subcellular localization studies of *Ti*SGD and *Ti*GS (2).

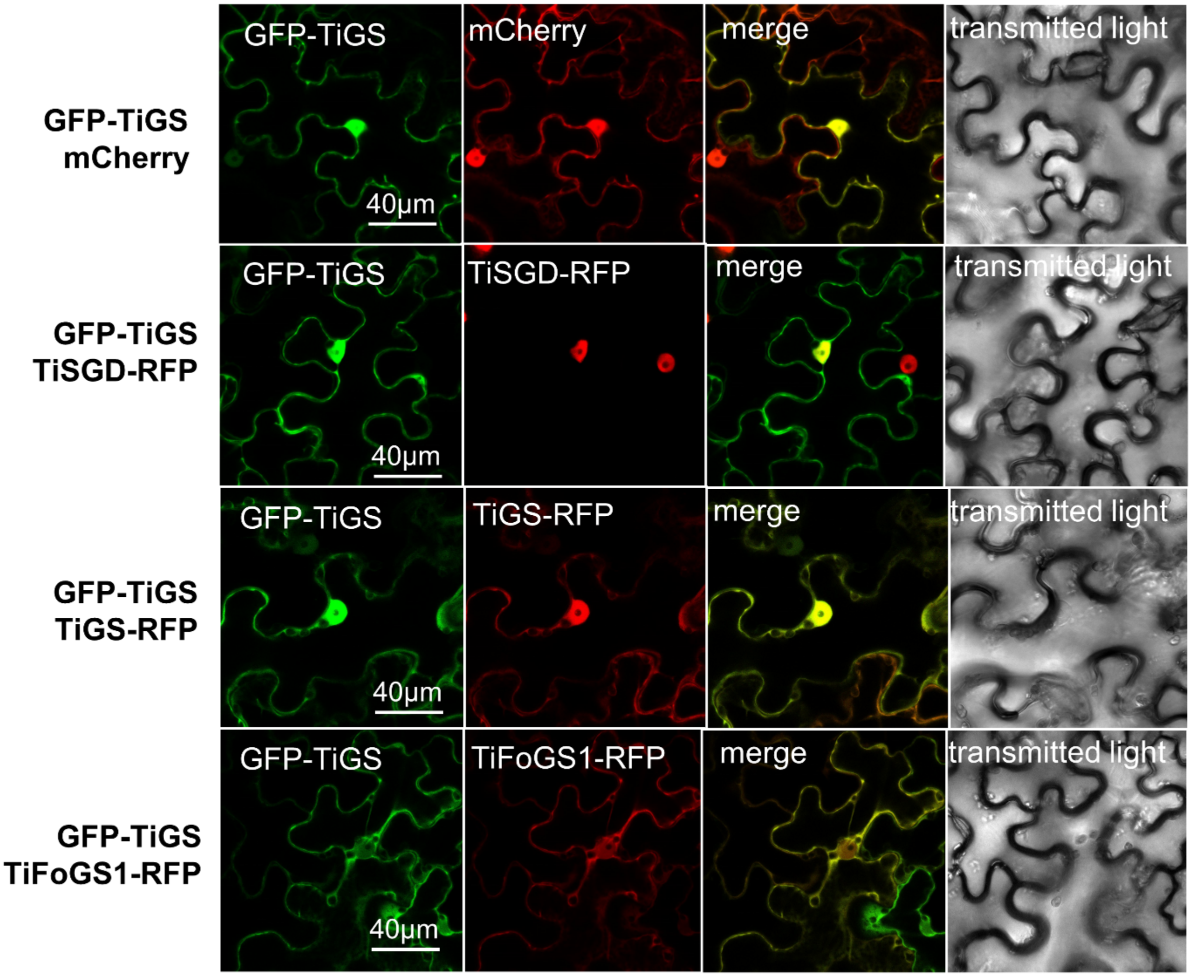


**Figure S17: Subcellular localization of *T. iboga* GS when co-expressed with SGD, GS, or FoGS1 in *N. benthamiana* leaves.** N-terminal GFP tagged *Ti*GS was co-expressed with C-terminal RFP tagged genes of interest. *Ti*GS co-expression with nucleocytosolic free-mCherry is shown for reference. GFP fluorescence is shown in green, RFP/mCherry in red, and merged GFP-RFP/mCherry in yellow.

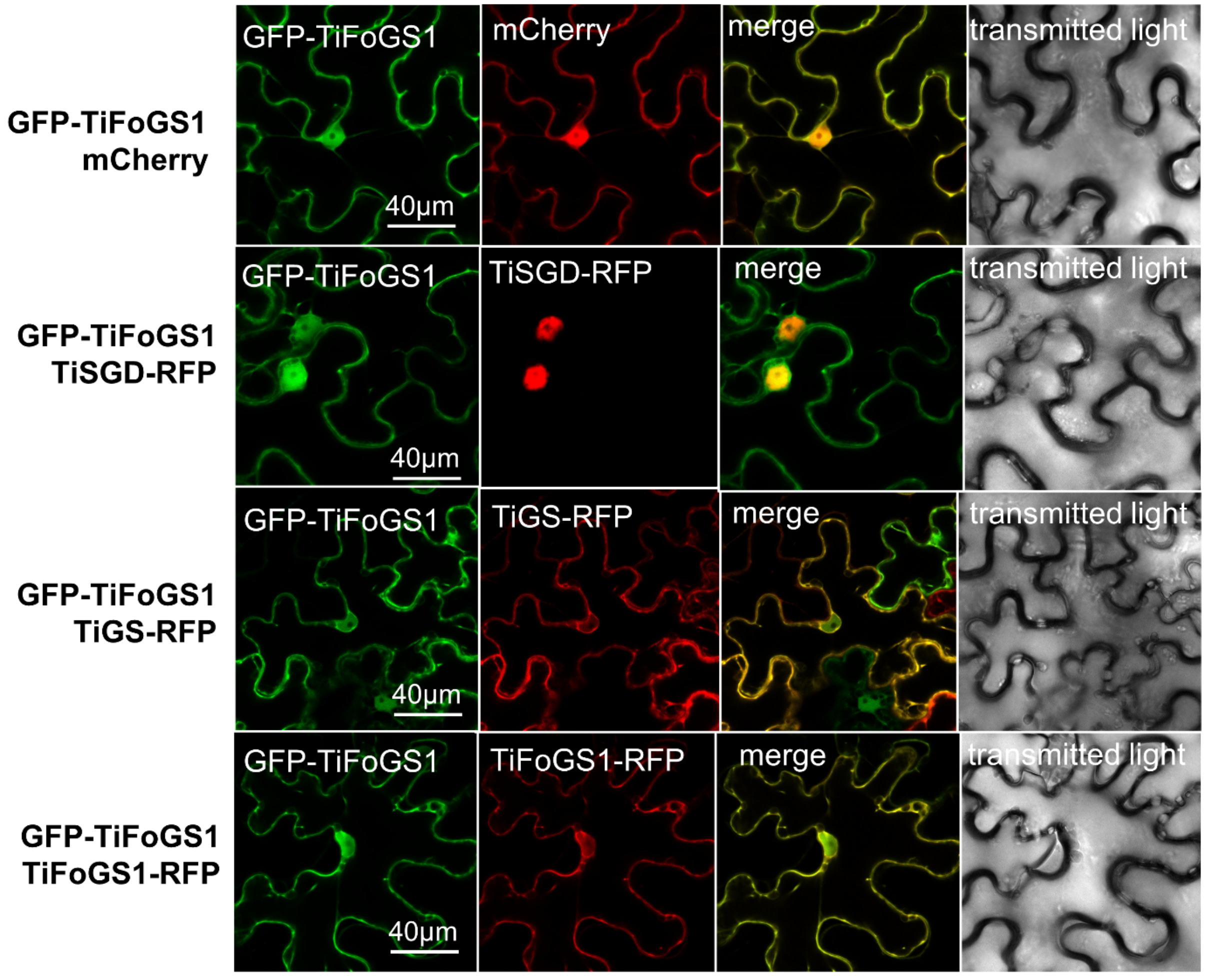


**Figure S18: Subcellular localization of *T. iboga* FoGS1 when co-expressed with SGD, GS, or FoGS1 in *N. benthamiana* leaves.** *Ti*FoGS1 with an N-terminal GFP tag was co-expressed with genes of interest with C-terminal RFP tags. *Ti*FoGS co-expression with nucleocytosolic free-mCherry is shown for reference. GFP fluorescence is shown in green, RFP/mCherry in red, and merged GFP-RFP/mCherry in yellow.

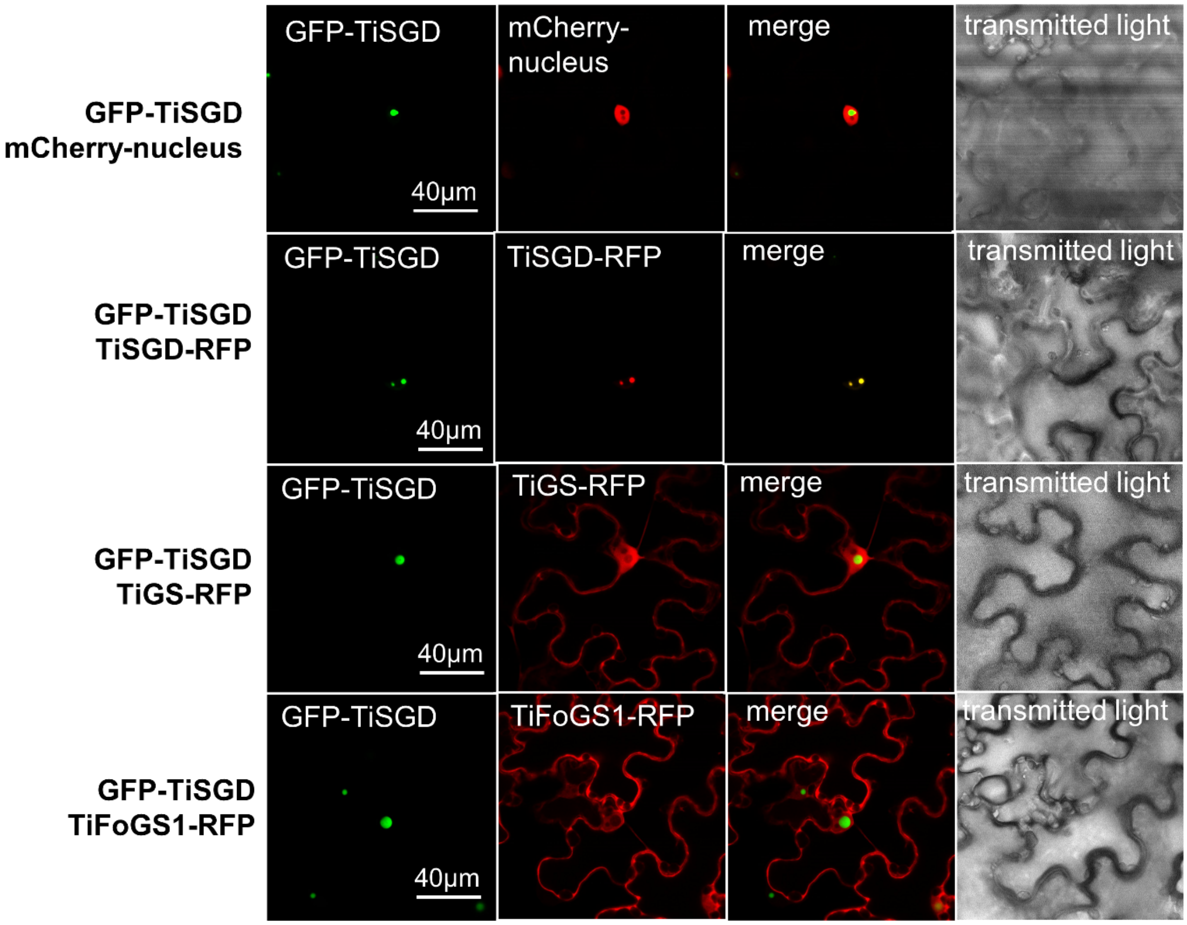


**Figure S19: Subcellular localization of *T. iboga* SGD when co-expressed with SGD, GS, or FoGS1 in *N. benthamiana* leaves.** N-terminal GFP tagged *Ti*SGD was co-expressed with C-terminal RFP tagged genes of interest. *Ti*SGD co-expression with nucleus-mCherry is shown for reference. GFP fluorescence is shown in green, RFP/mCherry in red, and merged GFP-RFP/mCherry in yellow.

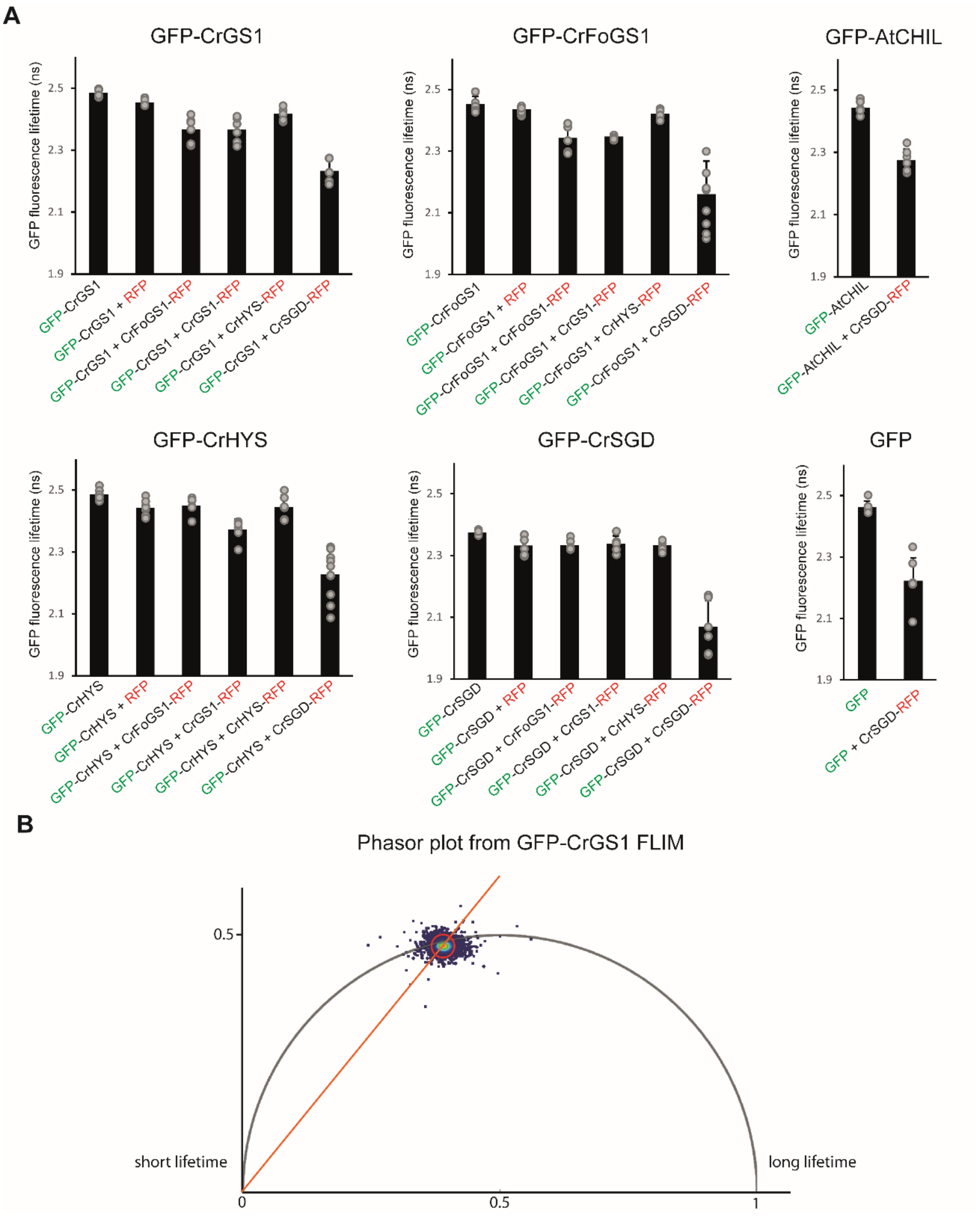


**Figure S20: GFP fluorescence lifetimes measured from FLIM-FRET assays of *C. roseus* SGD, FoGS1, GS1, and HYS in *N. benthamiana* leaves. A** Measured fluorescence lifetimes of GFP used to determine FRET in Figure 4B. Average GFP fluorescence lifetimes were measured from 6 biological replicates. All lifetimes were measured using phasor plot analysis (22). In instances where more than one GFP fluorescence lifetime was present in the same image, both were recorded. This only occurred when an image contained several nuclei with *Cr*SGD-RFP. Values for individual replicates are shown as grey circles and averages as black bars with error bars. We note the peculiar behavior of SGD-RFP, for which strong quenching of GFP was observed for all treatments, including a free-GFP negative control and *Arabidopsis thaliana* chalcone isomerase-like (CHIL). **B** A representative phasor plot of GFP-GS.

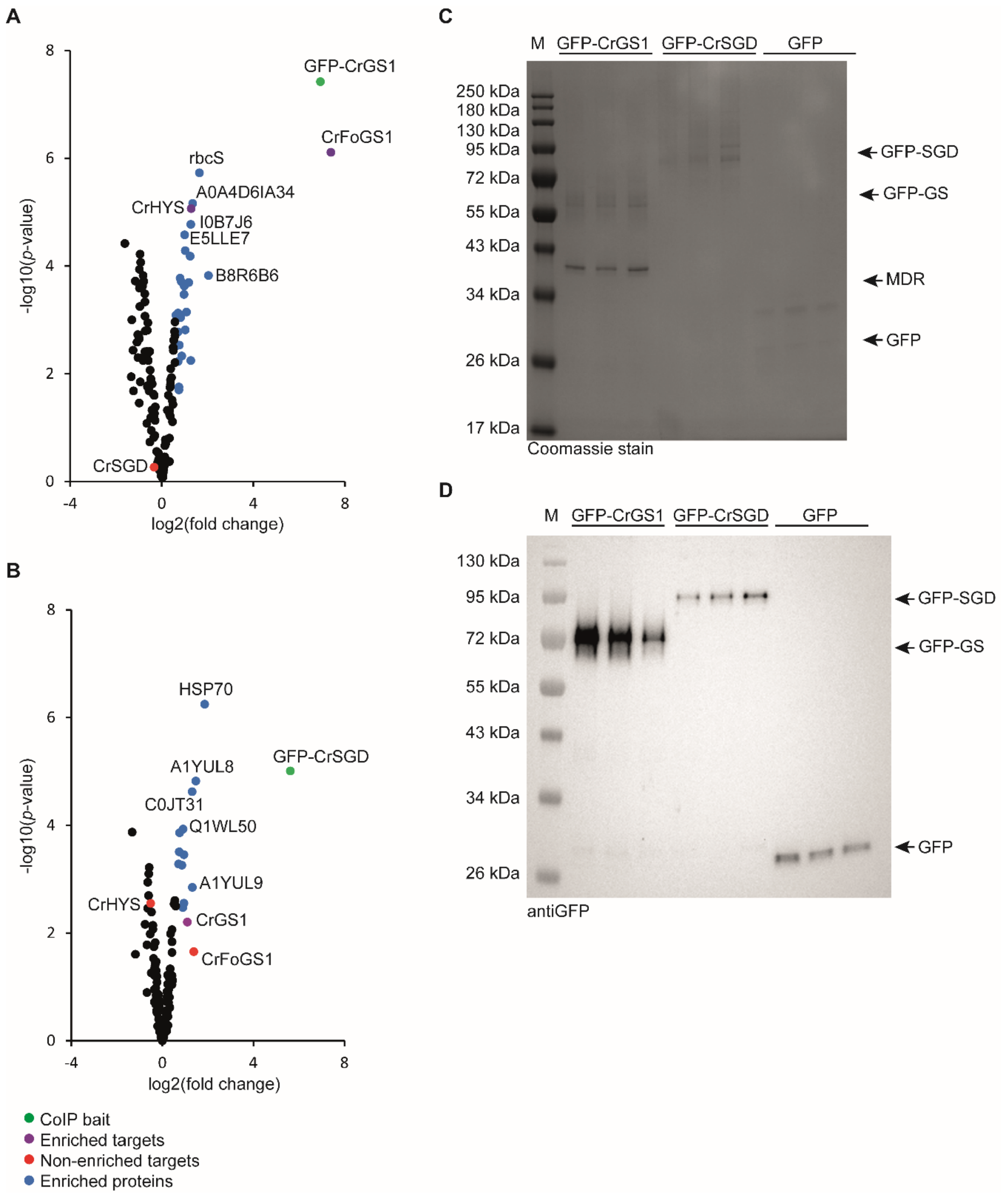


**Figure S21: Co-immunoprecipitations of GFP-*Cr*GS1 and GFP-*Cr*SGD. A** Co-immunoprecipitation of GFP-*Cr*GS1. **B** Co-immunoprecipitation of GFP-*Cr*SGD. GFP-tagged baits are shown in green, enriched proteins in blue, enriched target proteins in purple, and non-enriched target proteins in red. **E** SDS-PAGE analysis of co-immunoprecipitated proteins. M corresponds to the molecular ladder and Novex™ 12% gels were used. **D** Western Blot analysis of co-immunoprecipitated proteins using a GFP polyclonal antibody-horseradish peroxidase conjugate (Invitrogen) at a 1:1000 dilution followed by detection using Clarity Western ECL Substrate (*Bio-Rad*).

A

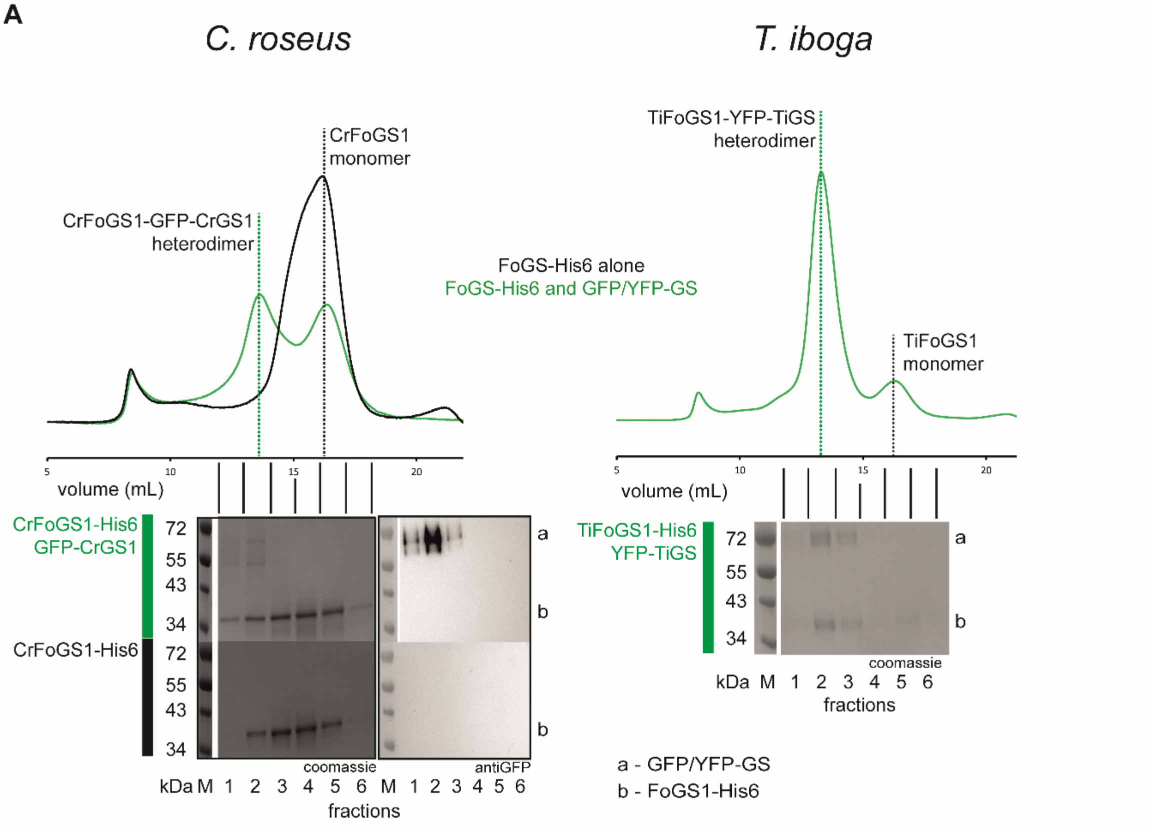


B

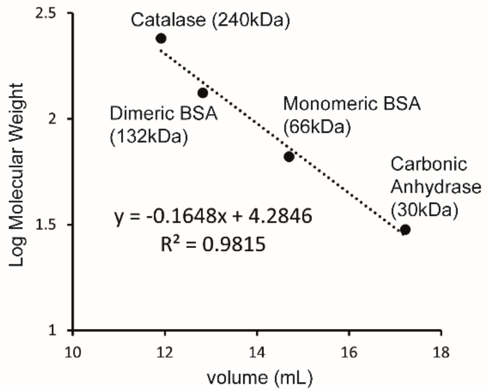


C

| Protein | Theoretical Weight (kDa) | Experimental Weight (kDa) | % Error |
|---|---|---|---|
| CrFoGS1 | 38.4 | 39.5 | 2.9 |
| GFP-CrGS1 | 69.5 | 83.6 | 20.3 |
| CrFoGS1-GFP-CrGS1 | 107.9 | 123.1 | 14.1 |
| TiFoGS1 | 38.8 | 39.2 | 1.0 |
| YFP-TiGS | 69.9 | 66.8 | 4.4 |
| TiFoGS1-GFP-TiGS | 108.7 | 106.0 | 2.4 |

**Figure S22: Size exclusion chromatography of purified *C. roseus* and *T. iboga* FoGS1-GS complexes.**

**A** Chromatograms from the size exclusion chromatography of co-purified *C. roseus* FoGS1 and GFP-GS1 and *T. iboga* FoGS1 and YFP-GS. SDS-PAGE of collected fractions are shown below each chromatogram with antiGFP Western Blot of GFP-*Cr*GS1 containing fractions. M corresponds to molecular ladders and Novex$^{TM}$ 12% gels were used. Green corresponds to co-purified FoGS1 and GFP/YFP-GS and black corresponds to FoGS1 purified without GFP-GS co-expression. **B** Standard curve used to calculate molecular weights. **C** Comparison of theoretical molecular weight determined using protparam (12) and experimental molecular weights calculated from elution volume. Percent error between theoretical and experimental values are shown. Experimental molecular weight of GFP/YFP-GS was calculated by subtracting the FoGS1 monomer weight from the FoGS1-GFP/YFP-GS heterodimer weight. BSA; bovine serum albumin.

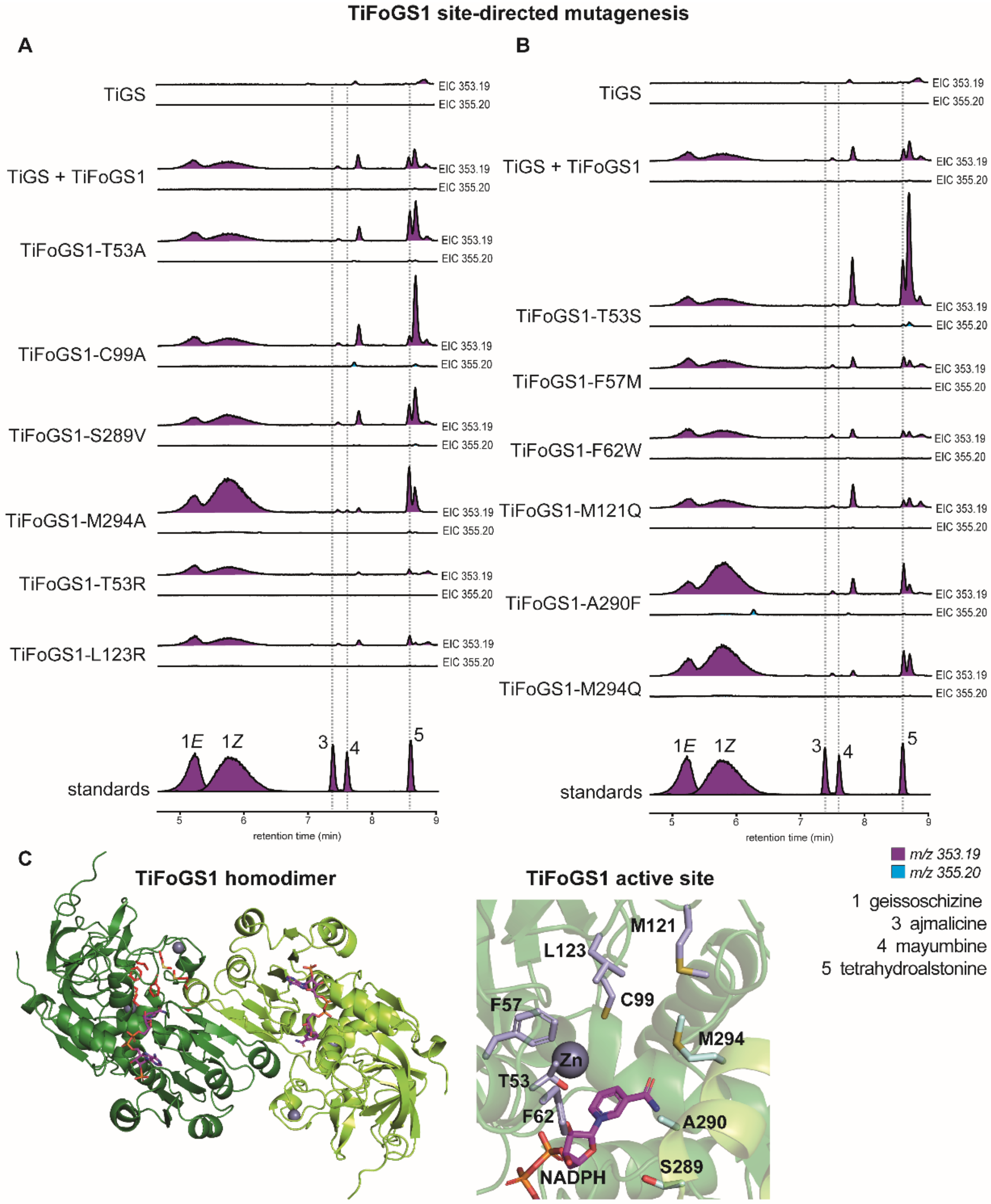


**Figure S23: Site-directed mutagenesis of *Ti*FoGS1.** *In vitro* GS-coupled assays of *Ti*FoGS1 variants. Extracted ion chromatograms (EIC) are shown for *m/z* values of *353.19* (purple) and *355.20* (cyan). Standards for 19*E*/*Z*-geissoschizine (1), ajmalicine (3), mayumbine (4), and tetrahydroalstonine (5) are shown below. **A** Site-directed mutagenesis targeting residues which could contribute to catalysis. The T53R and L123R variants were indented to introduce a large amino acid side chain into the *Ti*FoGS1 active site to disrupt non-GS activity. **B** Site-directed mutagenesis targeting non-conserved residues between *Ti*FoGS1 and *Ti*FoGS3 **C** Alphafold3 prediction (ipTM = 0.93; pTM = 0.94) of a *Ti*FoGS1 homodimer. Dark and light green distinguish protomers. NADPH is shown in magenta, targeted amino acids for site-directed mutagenesis in light purple or cyan (for residues contributed by dimerization), and the coordinated zinc ion as a grey sphere. Carbon atoms are coloured green/magenta, blue for nitrogen, red for oxygen, and yellow for sulfur.

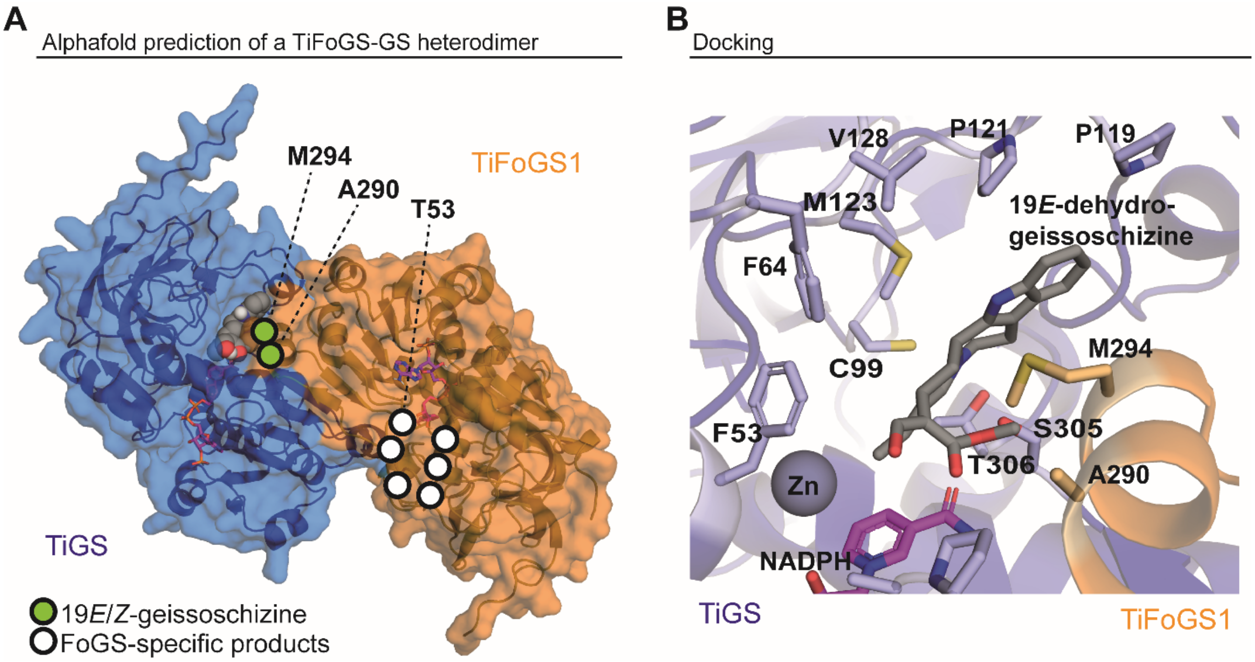


**Figure S24: A** Alphafold3 prediction (ipTM = 0.94; pTM = 0.93) of the *T. iboga* FoGS1-GS heterodimer. *Ti*GS is shown in blue, *Ti*FoGS1 in orange, NADPH in magenta, and docked 19*E*-dehydrogeissoschizine as grey spheres. The location of amino acids from site-directed mutagenesis of *Ti*FoGS1 and the corresponding effect of geissoschizine formation (green) and FoGS1- specific product formation (white) are mapped onto the model. **B** Molecular docking of 19*E*-dehydrogeissoschizine (grey) into the Alphafold3 prediction of the *T. iboga* FoGS1-GS heterodimer. The colouring scheme matches that of panel A. The coordinated zinc ion is shown as a dark grey sphere. Carbon atoms are shown in the base colour, nitrogen in blue, oxygen in red, and sulfur in yellow.

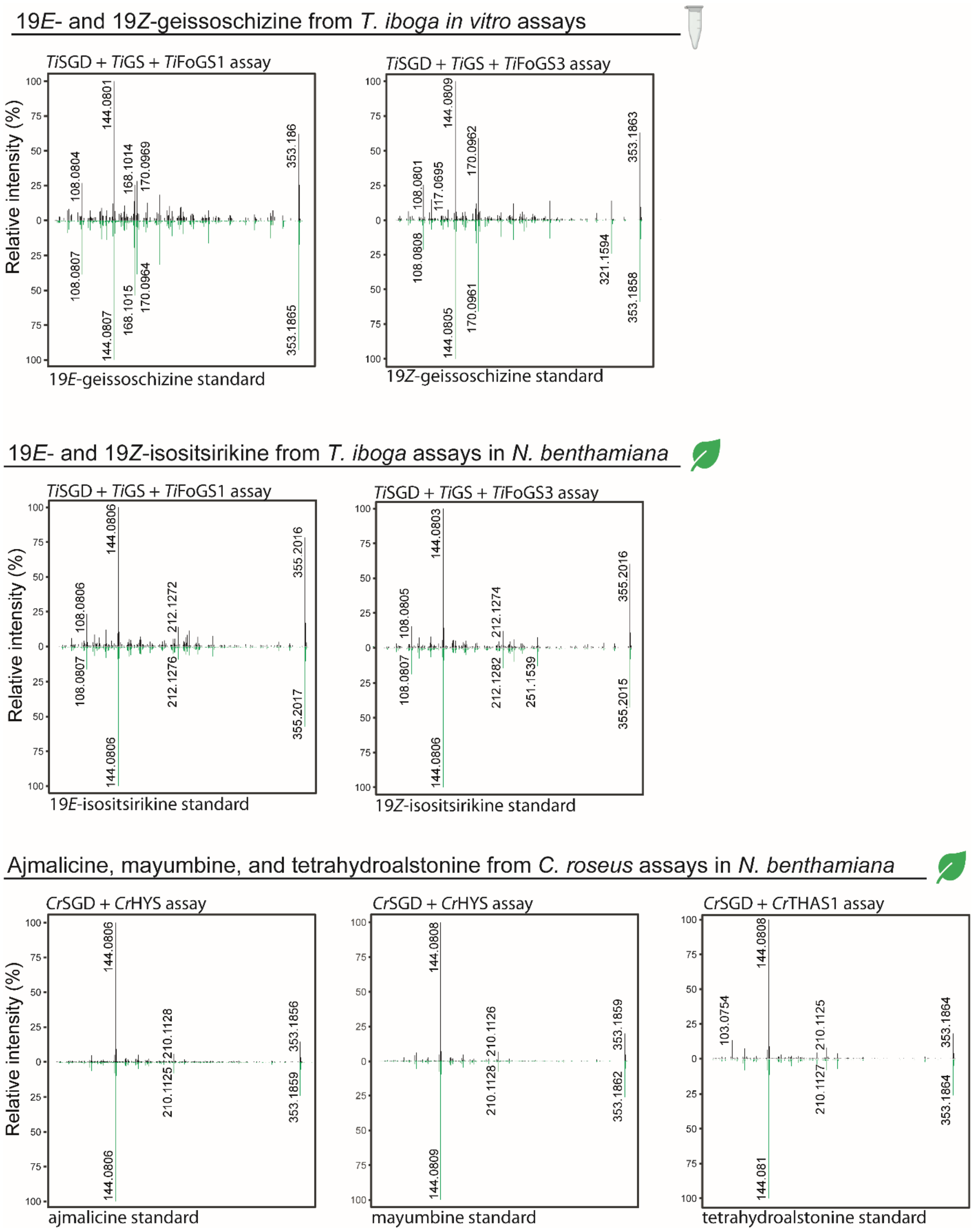


**Figure S25: MS/MS of analytes and authentic standards.** Representative MS/MS spectra for the major analytes of interest in this study compared to authentic standards: 19*E*-geissoschizine, 19*Z*-geissoschizine, 19*E*-isositsirikine, 19*Z*-isositsirikine, ajmalicine, mayumbine, and tetrahydroalstonine. MS/MS data for stemmadenine and catharanthine are shown in Figure S4 and S9, respectively.

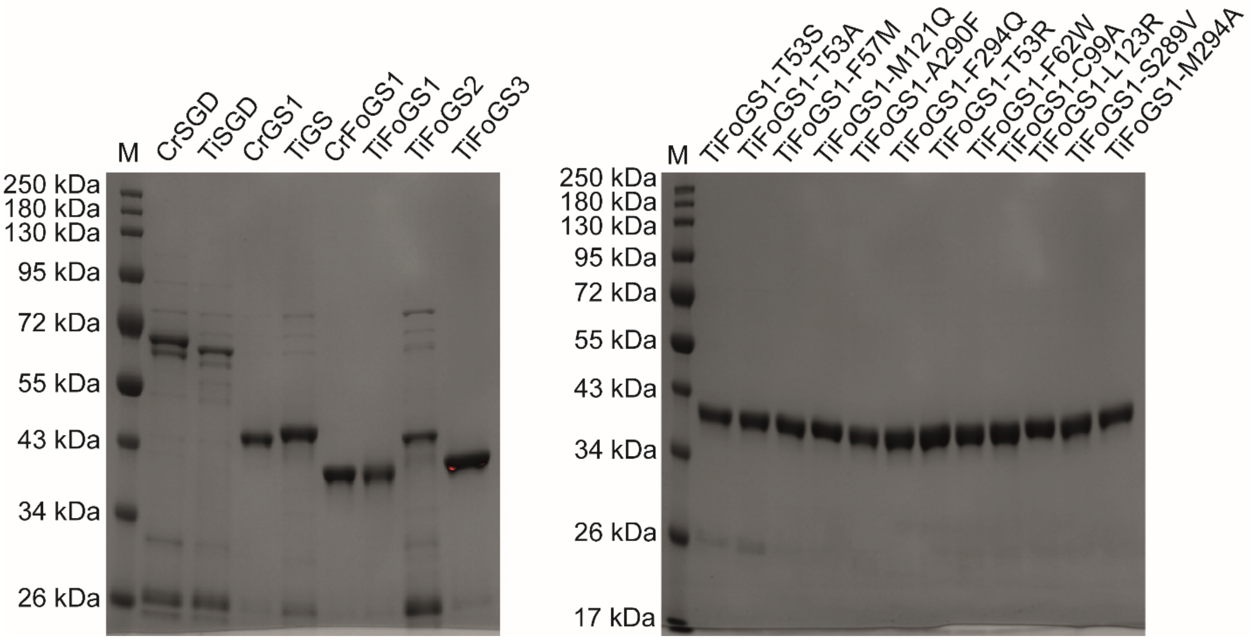


**Figure S26: SGS-PAGE of purified recombinant proteins.** M corresponds to molecular ladders and Novex™ 12% gels were used and stained with Coomassie.

### Supplemental Tables

#### Table S1: primers for amplification from *C. roseus* and *T. iboga* cDNA

| Gene name | Gene ID | GeneBank ID | Gene specific region of forward primer | Gene specific region of reverse primer |
|---|---|---|---|---|
| CrSGD | CRO-03G003560 | AF112888 | ATGGGATCGAAAGATGATCAG | TTAGTATTTTTGCTTCTTGACTAACTC |
| CrGS1 | CRO-06G024600 | KF302079 | ATGGCCGGAGAAACAACC | TTCCTCAAATTTCAATGTATTTCCAATG |
| CrHYS | CRO-01G032400 | KU865325 | ATGGCTGCAAAGTCACCTG | CTAGAAAGATGGGGATTTGAGAG |
| CrTHAS1 | CRO-01G033230 | KM524258 | ATGGCAATGGCTTCAAAGTC | CTAATTTGATTTCAGAGTGTTCCCTATAT C |
| CrFoGS1 | CRO-06G024580 | PZ105964 | ATGGCCGGAAAATCACCAG | AGGAGCTTTCAAGGTCTTTGC |
| CrFoGS2 | CRO-03G009850 | PZ105965 | ATGCCACCTCATTCCACATG | TTAAGGAGCTCTCAAGGTGTTG |
| CrFoGS3-like | CRO-06G024550 | PZ105966 | ATGGAGAAATCACCTGCAGAAAT | TTAAGCAGATTTCAATGTGTTGCC |
| CrADH9 | CRO-03G022230 | KF302077 | ATGGCCAGAAAATCACCAG | CACCTCTGATGGAAGAGTGAG |
| CrADH21 | CRO-08G001030 | KU865328 | ATGGCTCAAACAACTCCAAAC | AAGATTAGATGATTTGGAAGCTATATCG |
| Cr8HGOa | CRO-06G024560 | AY352047 | ATGGCGAAATCACCGGAAG | TTATGCAGATTTCAGTGTGTTGG |
| HL1 | CRO-08G004460 | PZ105975 | ATGCCGCCGGCTATGATTTC | TTAATCCGTTGAATTTTTCCAAATCTG |
| HL2* | CRO-06G018880 | NA* | ATGGTGCATCCGGTGGC | CTAGCCGTCCACCGAAC |
| HL3 | CRO-03G001660 | PZ105976 | ATGAGTTTTACTGGCCCTTCT | TTATACTTTTCCCTCAAGCCC |
| MLP1 | CRO-02G010700 | PZ105970 | ATGTCTGATCAATCTTTAGCTGG | TCACTTGAGATGGTGAGTCTC |
| MLP2 | CRO-02G026670 | PZ105971 | ATGGCTCCTCAAACTCTTGAT | TTAGATCTCATTGCCAGCAAGA |
| MLP3 | CRO-06G004160 | PZ105972 | ATGGCTTCTGTAGTTACTTTCAC | TTAAACATAGGCATCAGGATTTGC |
| MLP4 | CRO-02G016790 | PZ105973 | ATGGGTATCAAAGGGAAAATGATC | TCAAGAGTGGTGAGCTTCAATA |
| MLP5 | CRO-01G026840 | PZ105974 | ATGGGATTGATTGAAGTTGATATTGA | TTACTTTTGACCTTCTTCTTGAAGAT |
| MLP6* | CRO-02G016730 | NA* | ATGGGTCTCAAGGGGAAAATG | TCAAGAGTGGTGAGCTTCAAT |
| TiFoGS1 | Taibo.S173960 | PZ105967 | ATGGCTGGAAAATCACCAGAAG | TTAAGGAGCTTTCAAGGTGTTG |
| TiFoGS2 | Taibo.S075670 | PZ105968 | ATGCTGATATCTCCATCTGTACAC | TTAAGGAGCTTTCAAGGTGTTG |
| TiFoGS3 | Taibo.S174000 | PZ105969 | ATGGCGAAGTCACCAGAAG | CTAGGCAGACTTCAAGGAATT |

*Could not be amplified from cDNA

#### Table S2: Primers for *T.* iboga FoGS1 site-directed mutagenesis

| Variant | Forward primer | Reverse primer |
|---|---|---|
| Wild type | ATGGCTGGAAAATCACCAG | NA |
| T53S | GTGTGTCATTCGGATCTTCATTTCCTCAAGAATGAATTTG | AAGATCCGAATGACACACCCCACAATAAAGCACCTTG |
| T53A | GTGTGTCATGCGGATCTTCATTTCCTCAAGAATGAATTTG | AAGATCCGCATGACACACCCCACAATAAAGCACCTTG |
| F57M | GATCTTCATATGCTCAAGAATGAATTTGGCCTTTCTATATAC | CTTGAGCATATGAAGATCCGTATGACACACCCCA |
| M121Q | TGTCCCAAACAGGTGTTAACCTATTCTACTCCATATTTTGATG | TAACACCTGTTTGGGACAATAGTTCTCCAGATCTGCAG |
| A290F | CTTCACTCTTTTCCTTTGCTTATGGGGAGAAAGATGG | CAAAGGAAAAGAGTGAAGGTCAAGTGGCTCGGCT |
| M294Q | CCTTTGCTTCAGGGGAGAAAGATGGTTTCTGGAAGTAAC | TCTCCCCTGAAGCAAAGGAGCAGAGTGAAGGTCAAG |
| T53R | GTGTGTCATCGCGATCTTCATTTCCTCAAGAATGAATTTG | AAGATCGCGATGACACACCCCACAATAAAGCACCTTG |
| F62W | AAGAATGAATGGGGCCTTTCTATATACCCCCTTGTAC | AAGGCCCCATTCATTCTTGAGGAAATGAAGATCCGTA |
| C99A | GTTGGCGCGTTAGTTGGTTCATGCCGCACTTGTG | ACCAACTAACGCGCCAACTCCAACTTTGTCTCCAAC |
| L123R | ATGGTGCGCACCTATTCTACTCCATATTTTGATGGAAC | AGAATAGGTGCGCACCATTTTGGGACAATAGTTCTCC |
| S289V | GACCTTCACGTGGCTCCTTTGCTTATGGGGAGAAAGA | AGGAGCCACGTGAAGGTCAAGTGGCTCGGCTGG |

| | | |
|---|---|---|
| M294A | TTGCTTGCGGGGAGAAAGATGGTTTCTGGAAGTAACATT | CTTTCTCCCCGCAAGCAAAGGAGCAGAGTGAAGGT |
| Wild type | NA | TTAAGGAGCTTTCAAGGTGTTG |

**Table S3: *C. roseus* FoGS sequences**

| Gene name | Gene ID | Nucleotide sequence |
|---|---|---|
| *Cr*FoGS | CRO-06G024580 | ATGGCCGGAAAATCACCAGAAGAGGAGCACCCAGTCAAGACCTATGGATTGGCTGCTC<br>ATGATTCATCTGGGGTTTTATCTCCGTTCAAATTCTCCAGGAGGGCAACTCTTGAGGAT<br>GATGTGAGGTTCAAGGTGCTATATTGTGGGATTTGTCATACTGACCTTCATTTCGCTAAG<br>AATGAGTGGGGTATTTCGACCTATCCTCTTGTACCAGGACATGAAATCGTAGGGGAAGT<br>TACAGAGGTCGGCGGCAAAGTTACAAAGGTCAAGGTTGGAGATAAAGTTGGTGTTGGC<br>TGCTTGGTTGGTTCATGCCGCACTTGTGATAATTGTCGTGCAGATCTTGAGAACTATTGT<br>CCCAAAATGGTGCTAACCTATGCAAGTCCAAACGTTGATGGAACGATTACCTATGGAGG<br>CTATTCCAATGAGATGGTATGCAATGAACACTTTATTGTTCGTTTCCCAGAGAACCTACC<br>ACTTGATGGTGGGGCACCATTGCTTTGTGCCGGTATTACTGTGTACAGTCCAATGAAAT<br>ACTATGGCTTTGCCAAACCCGGGAGCCACATAGCTGTTAATGGTCTTGGTGGACTTGGC<br>CATGTGGCTGTTAAGTTTGCAAAGGCCATGGGAGCAAAAGTGACAGTTATAAGTACATC<br>TGAGGGCAAGAAAGACGATGCCCTCAATCGTTTGGGTGCAGATGCATTTTTGTTGAGCA<br>GTAATCCAGAAGCACTGCAGGCTGCAACAGGCACATTTGATGGCATACTTAATACTATTT<br>CTGCTAAGCACGCTATTATCCCATTGCTTGGTCTACTAAAGTCTCATGGCAAGCTTGTTC<br>TTCTTGGGGCACCCCCGGAACCACTTGATCTTCACTCTGCTCCTTTGCTTATGGGGAGG<br>AAGATGGTTGCTGGAAGTAGCATTGGAGGATTGAAGGAGACCCAAGAGATGCTTGATTT<br>TGCCGGAAAGCATAACATTACTGCAGATATAGAACTCATTTCCGCGGACAATATCAACA<br>CAGCTTTGGAGCGTCTGGCCAAGGGTGATGTTAGATATCGCTTTGTCCTTGACGTTGCA<br>AAGACCTTGAAAGCTCCTTAA |
| *Cr*FoGS2 | CRO-03G009850 | ATGCCACCTCATTCCACATGTACAATTGTGGGTAGATATTTTTTTTGCATGTTGATTTCTG<br>CCGTTTCACACCCAAAACCACCACTCTTCACCACAGCTCCACCGCCCGAACTACATTTC<br>GTGACCGCCGCCGGTAGGAAAATTTCAAAGATAATAAATATGGCGGGAAAATTACCAGA<br>ACAAGTTTTTCCGGTGAAGGCTCATGGATGGGCTGCTCATGACTCATCCGGAATTCTCT<br>CCCCTTTCAATTTCTCCAGAAGGGAGACGCTGGAGGATGATATCAGGTTCAAGGTTCTG<br>TATTGTGGTATTTGTCATACTGATCTTCATTTCATCAAGAACGATTGGGGCATATCAAGA<br>TATCCTCTTTTACCAGGACATGAAATTGTTGGCGAAGTTACAGAGGTGGGAAGCAAGGT<br>TACAAAAGTAAAAATTGGAGATAAAGTGGGGGTTGGCTACTTGGTTGGATCATGTCGTA<br>GTTGTGACAATTGTTCAGTAAATCTTGAGAACTATTGTCCCAAATCGGTTCTAACGGCTG<br>GAGCTGTTTATTTTGACGGCACCCCAACGTATGGTGGCTTTTCAAATGAAATGGTATGTA<br>ATGACAACTTTGTGGTTCGTTTCCCGGACACTTTGCCACTTGATGCTGGCGCTCCGTTG<br>CTTTGTGCCGGTGTCACCGTGTATAGTCCGATGAAATACTATGGCTTTGCGAAGCCGGG<br>AAACCATGTCGGAGTTAACGGACTTGGCGGCCTTGGTCATGTGGCTGTTAAGTTTGCAA<br>AGGCTTTTGGGGCAAAAGTGACAGTTATTAGTAGATCTTCTAAGAAGAAAGAGGAAGCT<br>ATTGATTATCTTGGTGCAGATGCATTCTTAGTCAGCCAAAATCCAGAAGAAATGAAGGCT<br>GCAGCAGGGACCATGGATGGTATTATCGATTGTGTCTCGGCTAAGCATCAATTAGTGCC<br>GTTACTTGGTCTACTCAAGTATCATGGAAAACTTGTTCTTGTTGGAGTTCCACCAGAACC<br>ACTTGACCTTCCAGCCGCTCCTTTGATTACTGGGAGGAAACTGATCGGAGGAAGCAATG<br>TGGGAGGACTAAAGGAGACTCAAGAGATGATTGATTTTGCTGCACAGCACAATATAACG<br>GCAGATGTGGAGGTTATTTCTATGGATTATGTAAATACAGCTATGGATCGTCTTGCTAAA<br>GGTGATGTTAGATATCGCTTTGTCATCGACATCGGCAACACCTTGAGAGCTCCTTAA |
| *Cr*FoGS3 | CRO-06G024550 | ATGGAGAAATCACCTGCAGAAATTGAGCATCCAGTCAAGGCCTTTGGATGGGCTTCTAA<br>AGACAATTCTGGAGTTCTCTCCCCCTTTGTTTTCTCCCGAAGGGCTACCGGAGAGCACG<br>ATGTGCAGTTCAAAGTTTTGTACTGTGGGATTTGCCCCTCCGATGTTGAAATGATTACCA<br>ACAAATGGGGTTTCATCAAGTATCCAGTTGTCCCTGGACATGAGATTGTGGGAGTGGTA<br>ACTGAAGTTGGAATCAAAGTGGAGAAATTCAAGATTGGTGACAAAGTGGGCGTAGGACA<br>CCTAGTAGAATCATGCCGAAAGTGTGATTTATGTTTTAAGGATCTTGAAAATTACTGTCTT<br>GGCAAGAGATTTGCACATATCACTAAATACGACGAGACAGGAATCATTAGATTCGGAGG<br>TTTTTCTGATGTAATGGTTGCCGATGAACATTTTGTGATTCGTTGGCCTGAGAATTTGCC<br>TATGGATATTGGTGCTCCTTTGCTTTGTGCTGGGATTACTACATATAGTCCATTGAAACA<br>TTTTGGACTTGATAAACCAGGAATCCATATTGGCATTGTTGGTCTTGGTGGTCTTGGCCA<br>TATAGCTGTGAAATTTGCCAAGGCTTTTGGTGCAAAAGTTACCGTGATTAGTACATCTGA<br>GTTTAAAAAACAGGAGGCTTTAGAGAAACTTGGTGCCGATGCTTTCTTGTTCAGTGCCA<br>ATCCCGAGCAGATGCAGGCTGCAGAATGGACTATGGATGGAATTATAGATACAGTATCA<br>GCAGTTCATCCAATGTCGCCCCTCGTCATGTTATTAAAGAATGATGGGAAGCTTATTATG<br>ATCGGTGCACCTAGAAAACCAATTGAACTTGAAGCGAAACCTGTCATCCTTGGGAGGAA<br>AATAGTAGCTGGAAGTGTTATTGGGGGCTTGAAAGAAACTCAAGAAATGATCGATTTTG<br>CAGCAAAGCACAACATTTTACCGGAAGTGGAGGTTATTCCAGTGGACTATATAAACAAT<br>GCAACCAACAGAATCTTGAAAGCTGATGTTAAATACAGATTTGTAGTTGACATTGGCAAC<br>ACATTGAAATCTGCTTAA |

*Discrepancy between the *Cr*FoGS2 sequenced cloned from cDNA and the corresponding gene id is shown in red

**Table S4: *T. iboga* FoGS sequences**

| Gene name | Gene ID | Nucleotide sequence |
|---|---|---|
| *Ti*FoGS | Taibo.S 173960 | ATGGCTGGAAAATCACCAGAAGAGGAGCACCCAGTGAAGACCTATGGATGGGCTGCTCGA<br>GATTCATCTGGGGTTCTTTCTCCGTTCAAATTCTCCAGGAGGGCAACACTTGAGGATGATGT<br>TAGATTCAAGGTGCTTTATTGTGGGGTGTGTCATACGGATCTTCATTTCCTCAAGAATGAATT<br>TGGCCTTTCTATATACCCCCTTGTACCAGGGCACGAAATCGTAGGTGAAGTTACAGAGGTT<br>GGCAGCAAAGTTACAAAAGTCAAGGTTGGAGACAAAGTTGGAGTTGGCTGCTTAGTTGGTT<br>CATGCCGCACTTGTGCTAACTGTTCTGCAGATCTGGAGAACTATTGTCCCAAAATGGTGTTA<br>ACCTATTCTACTCCATATTTTGATGGAACCATTACATATGGGGGCTATTCGAACGAGATGGT<br>CTGCAATGAGCACTTTATCATTCGTTTCCCAGAGAACATGCCACTTGCTGCTGGTGCTCCAT<br>TGCTGTGTGCTGGAATTACTGTCTACAGTCCAATGATATACTATGGCATTGCCAAACCAGGA<br>AAGCACATAGGTATTAACGGTCTTGGTGGGCTTGGCCATGTAGCTGTTAAGTTTGCAAAGG<br>CTTTGGGAGCAAAAGTGACAGTTATCAGTTCATCTGAGAGCAAGAGAGATGAAGCCATAAAA<br>CGTCTTGGTGCAGATGCATTTTTGGTGAGCACAAAGCCAGAAGAATTGCAGGCTGCAACAG<br>GCACAATGGATGGTGTACTTGATTGTGTTTCTGCTAAGCACCCAATTATCCCATTGCTTGGT<br>CTCCTCAAGTATCATGGAAAGCTTTGTATAGTTGGGGCACCAGCCGAGCCACTTGACCTTC<br>ACTCTGCTCCTTTGCTTATGGGGAGAAAGATGGTTTCTGGAAGTAACATTGGAGGATTGAAG<br>GAGACTCAAGAGATGATTGATTTTGCCGCAAAGCACAACATCACTGCAGATATTGAGATGGT<br>TTCAATGGACAACATCAACTTAGCCCTTGAGCGCCTTGCCAAGGGTGATGTTAGATATCGCT<br>TTGTCATCGACGTTGCCAACACCTTGAAAGCTCCTTAA |
| *Ti*FoGS2 | Taibo.S 075670 | ATGCTGATATCTCCATCTGTACACAGAAGACTGCCCCACTTCCCGACACCTCCGCCGGGAC<br>TACTATACGAGACCGTCCGTCGGAAGTTTACCTCGATAACAATGGCCGAAAAATCACCGGA<br>AGAGCTGTTCCCACTGAAGACTCATGGATGGGCTGCTCGTGACTCATCTGGGATTCTCTCC<br>CCTTTCAAATTCTCCAGGAGGGAGACCCTGGAGGATGATGTCAGATTCAAGGTTCTCTATTG<br>CGGTATCTGTCATACTGATCTTCACTTAATCAAGAATGATTGGGGCATATCTCTATATCCTCT<br>TTTACCAGGGCACGAGATTGTAGGTGAAGTTACAGAGGTTGGAAGCAAGGTTACAAAAGTA<br>AAAGTTGGAGATAAAGTGGGCGTTGGCTACTTGGTTGGATCATGCCGTAATTGTGATAATTG<br>TTCAGCAGACTTGGAGAACTATTGTCCCAAAACGATTCTCACAGCTGGATCCCGTTATTTTG<br>ATGGCTCCCCTACATATGGAGGCTTTTCGAATGAAATGGTATGCAACGAGCACTTTGTTGTT<br>CGTTTCCCGGACAACTTACCACTTGATGCCGGTGCTCCATTGCTTTGTGCCGGAGTCACTG<br>TCTACAGTCCAATGAAATACTATGGCTTTGCCAAACCGGGAAACCACGTTGGAGTTAACGG<br>CCTTGGCGGGCTTGGTCACGTGGCTGTTAAGTTTGCAAAGGCCATGGGAGCAAAAGTGACA<br>GTTATCAGTAGATCTTCTAATAAGAAGGAGGAAGCTATAGATCATCTTGGCGCAGATGCATA<br>CTTGGTGAGCCAAGATCCAGAAGAAATGAAGGCTGCGGCAGGCACCATGGATGGTATAATT<br>GATTGTGTCTCGGCTAAGCACCAATTGCTGCCATTAATTGCTCTACTCAAGTATCATGGAAA<br>AATTGTTCTGGTTGGGGTACCAGCAGAGCCACTTGAGCTTTCAGCTGGTTCTCTGATTCTTG<br>GAAGGAAGATGGTTGCTGGAAGTAGTGTCGGAGGATTAAAGGAGACTCAGGAGATGATTGA<br>TTTTGCTGCAAAACACAGCATCACTGCAGATGTCGAGGTTATTCCCATGGACTATGTCAACA<br>CAGCTATGGAGCGTCTTGCTAAAGGTGATGTTAGATACCGCTTTGTGATCGACATTGGCAAC<br>ACCTTGAAAGCTCCTTAA |
| *Ti*FoGS3 | Taibo.S 174000 | ATGGCGAAGTCACCAGAAGTTGAGCATCCAGTTAAGGCCTATGGCTGGGCTGCTAGAGACC<br>TCTCCGGAGTTCTCTCCCCCTTCAATTTTTCACGAAGGGCTACGGGAGAGCATGATGTGCA<br>GTTCAAGGTGTTGTACTGCGGGATCTGTCACTCGGATCTTCACATGATTAAGAATGAATGGG<br>GAATCACCAAGTATCCTATTGTACCTGGGCATGAGATTGTTGGTGTGGTTACTGAGGTTGGC<br>AGCAAGGTGGATAAATTCAAGGTTGGGGATAAAGTAGGTGTAGGTTGCCTCGTTGGATCAT<br>GCCGCAAATGTGATATGTGCACCAAGGATCTTGAAAATTACTGTCCCGGTCAGATACTCACA<br>TATAGTGCTACCTACACTGATGGAAGCACTACATATGGAGGCTACTCTGATCTCATGGTTGC<br>GGATGAACACTTTGTGATTCGTTGGCCTGAAAATCTGCCTATGGAGACTGGTGCTCCTCTGC<br>TCTGTGCTGGGATCGCAACATACAGTCCATTGAGATATTTTGGACTTGACAAACCTGGAACC<br>CATGTTGGCATTGTTGGTCTTGGTGGTCTTGGCCATATGGGTGTGAAGTTTGCCAAGGCTTT<br>TGGGGCAAAAGTAACTGTCATAAGCACATCTGACAGTAAAAAGCAGGAAGCCATAGAGAAA<br>CTTGGTGCCGACGCATTCCTGGTCAGCCGTGATCCCCAGCAAATGCAGGCAGCATGTGGC<br>ACTTTGGATGGCATTATTGATACTGTGTCTGCACCTCATGCTCTTATGCCTCTGGTGAGCTT<br>ATTGAAGTCTCACGGAAAGCTTATAATGGTTGGTGCACCTGAAAAGCCACTTGAGCTGCCA<br>GTGTTTCCTCTCCTCCAAGGGAGGAAAATAATAGCCGGAAGTGCCATTGGAGGCTTGAAGG<br>AAACTCAAGAAATGATCGATTTTGCAGCAAAGCACCACATTTTACCTGATGTCGAGCTCATT<br>CGGATGGACTATGTTAACACTGCAATGGAGCGGGTCCTGAAATCCGATGTCAAGTACAGAT<br>TTGTCATCGACATTGGCAATTCCTTGAAGTCTGCCTAG |

**Table S5: Primers for amplification of the *Cr*FoGS1 VIGS target region**

| Gene | Forward primer | Reverse primer |
|---|---|---|
| *Cr*FoGS1 | ATATTGCTGCGGATCCCAGTAATCCAGAAGCACTGC | ATGCCCGGGCCTCGAGGTGTTGATATTGTCCGCGG |
